# Detecting Diffusion Imaging constructive connectivity analysis for “What” stream Visual Pathways with correlations to Visual Agnosia

**DOI:** 10.1101/486639

**Authors:** Ganesh Elumalai, Geethanjali Vinodhanand, Valencia Lasandra Camoya Brown, Jessica Dasari, Venkata Hari Krishna Kurra, Kevin Andrew Douglas, Nitya Akarsha Surya Venkata Ghanta, Kammila Joshna Naga Sudha, Divya Singh, Nitisha Tricia Dyal, Pravitha Jose Sherly, Ann Maria Joy

## Abstract

Visual perception is the ability to interpret the surrounding environment. A hypothetical “ventral” stream visual pathway explains how we perceive objects with respect to spatial orientation. The ventral stream fibers extend between “Visual cortex to Inferior temporal gyrus. The previous studies have failed to prove any indications on the -structural connectivity of this pathway. This study is designed to trace the existence of neural structural connectivity between “Visual cortex with Inferior temporal gyrus” using “Diffusion Tensor Imaging Tractography”, which aims to correlate its functional importance with visual object perception.

The observational analysis used thirty-two healthy adults, ultra-high b-value and diffusion MRI datasets from an Open access research platform. The datasets range from both sexes, between 20–49 years, with mean age of 30.4 years. The confirmatory observational analysis process includes; datasets acquisition, pre-processing, processing, reconstruction, fiber tractography and analysis using software tools. All the datasets confirmed the fibre structural extension between, “Visual cortex to Inferior temporal gyrus in both the sexes may responsible for the visual perception of objects. This new fiber connectivity evidence justifies the structural relevance of visual perception impairments, such as visual object agnosia.

## 1. INTRODUCTION

Past evidences which were proposed by most of the scientists that have published literature on ventral stream visual pathway states that it is a multisynaptic cortico-cortical connection. Ventral stream visual pathway is a cortical interconnection between the primary visual cortex (Brodmann area 17,18,19) and the inferior temporal lobe (Brodmann area 20), that helps in the identification of objects, colour interpretation, and face identification and recognition as mentioned by Mishkin and Ungerleider (1983) [1]. The three projections of the occipitotemporal interconnections are namely occipitotemporal-medialtemporal, occipitotemporal-orbitofrontal, and occipitotemporal-ventrolateral prefrontal pathways that are associated with exposure to long-term memory and the association of object-reward respectively (1990) [3]. Therefore, any damage to the inferior temporal lobe can lead to lack of recognition and memory to objects (1983) [1]. Similarly, in monkeys it is connected between striate, pre-striate and inferior temporal cortex (1983) [2]. Kravitz and Saleem stated that the occipitotemporal network which is especially for the ventral stream visual pathway is helpful for attentional, contextual, and feedback effects. There are six different cortical and subcortical systems where each system works on its own specialized behavioral, cognitive, or effective function. These six cortical systems collectively provide the most important purpose for the ventral stream visual pathway.

In Subcortical pathways, firstly: the unidirectional occipitotemporo-neostriatal pathway, second: the occipitotemporal-ventral striatum pathway and third: the occipitotemporal-amygdaloid pathway, whilst in cortical pathways, firstly: the occipitotemporal-medial temporal pathway, second: the occipitotemporal-orbitofrontal pathway and third: the occipitotemporal-ventrolateral prefrontal pathway. Anatomically, the ventral pathway is an internetwork for feed-forward and feedback projections which are unidirectional. Thus, it plays a major role in the perceptual identification, recognition of objects, spatial association, grasping and molding objects with the hands (1990) [3]. Some studies mention that there is white matter connection between the dorsal and ventral stream pathway (2015) [7]. Moreover, Polanen and Davere mentioned that the ventral stream receives information from the dorsal stream visual pathway and therefore refined the interpretation towards object’s representation (2015) [8]. Additionally, Goodale and Milner proposed that the “what” pathway enables us to identify and grasp objects spatially oriented (1992) [4]. Severe bilateral damage to the occipitotemporal pathway that connects from the primary visual cortex to the inferior temporal lobe is in association with the “what” pathway hence can lead to Apperceptive agnosia which is the inability to recognize an object due to failure of perception. Apperceptive visual agnosia entails deficit of perceptual recognition of objects, differentiation of colors, faces, hand movements and gestures (2003) [9]. Apperceptive visual agnosia is mainly associated with damage to the association between the occipitotemporal interconnection that prevents the individual from identifying and recognizing objects and also framing their hand accordingly to hold or pick an object (1994) [5]. In the case of Alzheimer’s in relation to visual agnosia, the damage to the occipitotemporal cortex leads to the absence of coordination and identification of objects thus, leading to a severe damage to the ventral stream visual pathway. (1999) [6].

## 2. Materials and Methods

The present study, designed to use the open access, ultra-high b-value, diffusion imaging datasets from Massachusetts General Hospital – US Consortium Human Connectome Project (MGH-USC HCP), which acquired from Thirty-two healthy adult participants (16 Males and 16 Females, between the ages of 20–49 years). The imaging data is available to those who register and agree to the Open Access Data Use Terms.

### 2.1. Participants

Data was acquired from thirty-two healthy adults between the ages of 20 to 59. A written consent was given by every participant and the procedure was carried out in accordance to the review board of their institution (MGH/HST) approval and procedures. All participants involved in this study, as the MGH-USC Adult Diffusion Dataset were scanned on the 3T CONNECTOM MRI scanner (see (Setsompop et al., 2013) for an overview) were housed at the Athinoula A. Martinos Center for Biomedical Imaging at MGH. A custom-made 64-channel phased array head coil was used for signal reception (Keil et al., 2013). No data regarding the thirty-two adults were saved in any other imaging modalities. Thirty-two healthy adults participated in this study (16 Females, 16 males, 20–49 years old; mean age = 30.4 years old). A written consent was given by every participant and the procedure was carried out in accordance to the review board of Partners Healthcare by their institution (MGH/HST) approval and procedures. Participants’ gender and age are available in the data sharing repository. Due to the limited sample size, there are some ages which we had only one participant. Given de-identification considerations, age information is provided in 5-year age bins.

**(Fig – 1).**
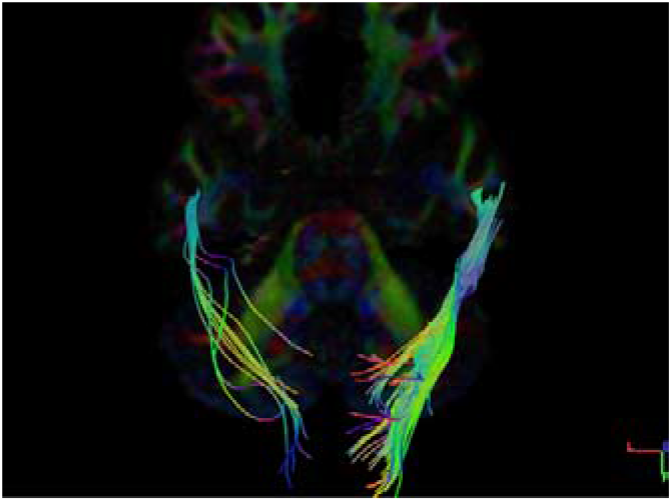
Female Bilateral Orientation of fibres

**(Fig – 2).**
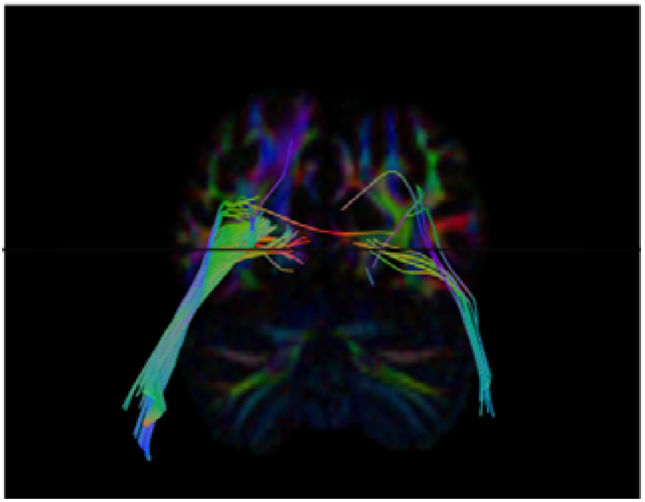
Male Bilateral Orientation of fibres

### 2.2 Data Acquisition

The data was collected on the customized Siemens 3T Connectome scanner, which is a modified 3T Skyra system (MAGNETOM Skyra Siemens Healthcare), housed at the MGH/HST Athinoula A. Martinos Center for Biomedical Imaging. A 64-channel, tight-fitting brain array coil was used for data acquisition. dMRI data was acquired using a mono-polar Stejskal-Tanner pulsed gradient spin-echo echo planar imaging (EPI) sequence with Parallel imaging using Generalized Auto calibrating Partially Parallel Acquisition (GRAPPA). The Fast Low-angle Excitation Echo-planar Technique (FLEET) (Polimeni et al., 2015) was used for Auto-Calibration Signal (ACS) acquisitions to reduce motion sensitivity of the training data and improve stability and SNR of the GRAPPA reconstructions. The Simultaneous Multi-Slice (SMS; multi-band) technique (Feinberg et al., 2010; Feinberg and Setsompop, 2013; Setsompop et al., 2012a; Setsompop et al., 2012b) has been shown to increase the time efficiency of diffusion imaging and was considered for this protocol, however at the point in the project timeline when data acquisition was to begin, the SMS method was not implemented in the EPI sequence featuring FLEET-ACS for GRAPPA. Because of the key benefits of in-plane acceleration for improved EPI data quality, such as lower effective echo spacing and hence mitigated EPI distortions and blurring, and also because of the longer image reconstruction times associated with SMS-EPI data, here data acquisition was performed without the use of SMS in favor of in-plane acceleration using FLEET-ACS and GRAPPA. (Acquisition of subsequent datasets including the Life Span dataset utilized a more newly implemented sequence combining SMS-EPI with FLEET-ACS and GRAPPA, see below.)

In each subject, dMRI data were collected with 4 different b-values (i.e., 4 shells): 1000 s/mm^2^ (64 directions), 3000 s/mm^2^ (64 directions), 5000 s/mm^2^ (128 directions), and 10000 s/mm^2^ (256 directions). Different *b*-values were achieved by varying the diffusion gradient amplitudes, while the gradient pulse duration (▯) and diffusion time (▯) were held constant. These *b*-values were chosen to provide some overlap with the HCP data from WU-Minn consortium, while at the same time push to the limit of diffusion weighting within the constraints of SNR and acquisition time. On determining the number of directions in each shell, in general, data with higher *b*-value was acquired with more DW directions to capture the increased ratio of high angular frequency components in the MR signal, and to compensate for the SNR loss due to increased diffusion weighting. Also, the number of directions in each shell was selected to meet the typical requirements of popular single shell and multi-shell analysis methods (e.g. *q*-ball imaging (Descoteaux et al., 2007; Tuch, 2004; Tuch et al., 2002; Tuch et al., 2003), spherical deconvolution (Anderson, 2005; Dell’Acqua et al., 2007; Tournier et al., 2004), ball and stick model (Behrens et al., 2007; Jbabdi et al., 2012), multi-shell *q*-ball imaging (Aganj et al., 2010; Yeh et al., 2010), diffusion propagator imaging (Descoteaux et al., 2009, 2011), etc.), and to keep the total acquisition time feasible in healthy control subjects.

The diffusion sensitizing direction sets were specifically designed so that the 64-direction set is a subset of the 128-direction set, which is again a subset of the 256-direction set. The initial 64 directions were calculated with the electro-static repulsion method (Caruyer et al., 2013; Jones et al., 1999). With these 64 directions fixed, another 64 directions were added as unknowns, and an optimized 128-direction set was calculated by adjusting the added 64 directions using electrostatic repulsion optimization. With these 128 directions fixed, the 256-direction set was generated using the same method. As such, all 4 shells share the same 64 directions; the b=5000 s/mm^2^ shell and the b = 10000 s/mm^2^ shell share the same 128 directions.

During data acquisition, the diffusion sensitizing directions with approximately opposite polarities were played in pairs to counter-balance the eddy current effects induced by switching the diffusion weighting gradient on and off. Each run started with acquiring a non-DW image (*b*=0), and one non-DW image was collected every 13 DW images thereafter. Therefore 552 image volumes were collected in total, including 512 DW and 40 non-DW volumes for each subject.

### 2.3 DTI Data Pre-processing and Quality Control

The MGH-USC HCP team completed their basic imaging data preprocessing, with software tools in Free surfer and FSL, which included (i.) Gradient nonlinearity correction, (ii.) Motion correction, (iii.) Eddy current correction, iv. b-vectors. All DTI data was corrected for gradient nonlinearity distortions offline (Glasser et al., 2013; Jovicich et al., 2006). Diffusion data was further corrected for head motion and eddy current artifacts. Specifically, the *b*=0 images interspersed throughout the diffusion scans were used to estimate bulk head movements with respect to the initial time point (the first *b* = 0 image), where rigid transformation was calculated using the boundary-based registration tool in the Free Surfer package V5.3.0 (Greve and Fischl, 2009). For each *b* = 0 image, this transformation was then applied to itself and the following 13 diffusion weighted images to correct for head motion. Data of all 4 *b*-values were concatenated (552 image volumes in total) and passed into the EDDY tool (Andersson et al., 2012) for eddy current distortion correction and residual head motion estimates. The rigid rotational components of the motion estimates were then used to adjust the diffusion gradient table for later data reconstruction purposes.

Both unprocessed and minimally pre-processed data are available for download in compressed NIfTI format. The measured gradient field nonlinearity coefficients are protected by Siemens as proprietary information. Because the gradient nonlinearity correction cannot be performed without this information, the unprocessed data provided had already been corrected for gradient nonlinearity distortions with no other pre-processing performed.

All the anatomical scans (T1w and T2w) were free from gross brain abnormalities as determined by a MGH trained physician. All MRI data (T1w, T2w and DW) went through the process of quality assessment by a MGH faculty member, MGH postdoctoral researcher and a MGH research assistant, all trained in neuroimaging in MGH. More specifically, each dataset was assessed by two raters, who viewed both the unprocessed and minimally pre-processed data volume by volume, and rated in terms of head movements, facial and ear mask coverage (to make sure that brain tissue was not masked off), and eddy current correction results. Finally, a comprehensive grade was given to determine whether a dataset had passed quality control.

### 2.4 SUMMARY OF DATASETS

This data set was originally from the MGH-USC HCP team which released *<u>diffusion imaging</u>* and structural imaging data acquired from 32 young adults using the customized MGH Siemens 3T Connectome scanner, which has 300 mT/m maximum gradient strength for diffusion imaging. A multishell diffusion scheme was used, and the b-values were 1000, 3000, 5000 and 9950 s/mm2. The number of diffusion sampling directions were 64, 64, 128, and 256. The in-plane resolution was 1.5 mm. The slice thickness was 1.5 mm.

### 2.5 FIBER TRACTOGRAPHY

Fiber tractography is a very elegant method, which can used to delineate individual fiber tracts from diffusion images. The main process of study, uses the “DSI-Studio” software tools for i. Complete preprocessing, ii. Fiber tracking and iii. Analysis.

The imaging data processing, helps to convert the pre-processed raw data to .src file format, which will be suitable for the further reconstruction process. The reconstruction of .src files achieved through a software tool. It converts the .src imaging data to .fib file format. Only the .fib files are compatible for fiber tracking. To delineate individual fiber tracts from reconstructed diffusion images (.fib file), using a DSI studio software tools.

## 3. RESULTS

The fibres were traced, and identified in all 32 Diffusion MRI datasets for “Ventral” stream pathways, involved in visual spatial perception. Considering the reconstruction of fibres originated from the Primary Visual Cortex, a confirmatory observation of neural structural connectivity to the inferior temporal lobule was traced using the software DSI studio. All Diffusion MRI datasets are derived from the Human Connectome project and were analysed in a repeated manner with provided effects and tools in the option window in the software. Observations on 32 MRI datasets, taking the age and sex into consideration the track analysis report retrieved. We performed t-test analysis for all 32 datasets selecting equal number of samples from both the group (16 males and 16 females) to check any difference in structural connections between the “Visual cortex (BA 18 & 19) to Inferior Temporal Lobule (BA 20)”. To validate the built atlas and its application among all the male and female datasets, we performed the t-test analysis of variance the for the following parameters (i) the number of tracts, (ii) volume of tracts, (iii) mean length of tracts and (iv) tracts standard deviation.

### i) Number of Tracts

The purpose of our study is deemed to find the neural structures extending between two cortices, connecting them from one end to another end. Fibre tracking uses diffusion tensor to track the fibres along the whole length starting from the seed to end region. The resolution of finding number of tracts helps us to correlate the actual function of ventral stream visual pathway – visual spatial perception with our present study. The greater number of fibres denotes that the patient has more efficiency in visually perceiving an object and a smaller number of tracts denotes that the patient might have less efficiency in perceiving an object or even a damage to that pathway may lead us to correlate clinically that the person might suffer from visual agnosia.

a. Male Right and Left Side
b. Female Right and Left side
c. Right side Male and Female
d. Left side Male and Female

#### a) Male Subjects Right & Lett

The Fibres were traced and observed for neural structural connectivity on both right and left side in all sixteen healthy adult male datasets **(Table 1).**

**Table 1:** Number of Tracts Right and left in Male *

| Subject | Number of Tracts Right | Number of tracts Left |
| --- | --- | --- |
| 1006 | 304 | 1 |
| 1007 | 47 | 6 |
| 1008 | 32 | 133 |
| 1009 | 12 | 21 |
| 1011 | 80 | 11 |
| 1013 | 243 | 17 |
| 1014 | 25 | 1 |
| 1015 | 3 | 1 |
| 1016 | 10 | 2 |
| 1022 | 3 | 55 |
| 1025 | 78 | 48 |
| 1027 | 221 | 272 |
| 1028 | 4 | 153 |
| 1029 | 367 | 54 |
| 1030 | 9 | 45 |
| 1031 | 105 | 45 |

\* $p$ -value is .236563, result is *not* significant at $p < .10$

Dataset numbers 1006, 1007, 1011, 1013, 1014, 1016, 1027, 1029, 1031 shows greater number of tracts at right side. Dataset numbers 1008, 1009, 1015, 1022, 1027, 1028, 1030 shows a greater number of tracts on left side. Therefore, from the table given below, 9 datasets have more fibers on right and 7 datasets have more fibers on left but the variation in the count of fibers are more on right side **(dataset number 1029).**

When the results were statistically compared by performing independent t – test no significant difference were seen between the number of tracts in right and left side of female subjects.

**(Figure-1).**
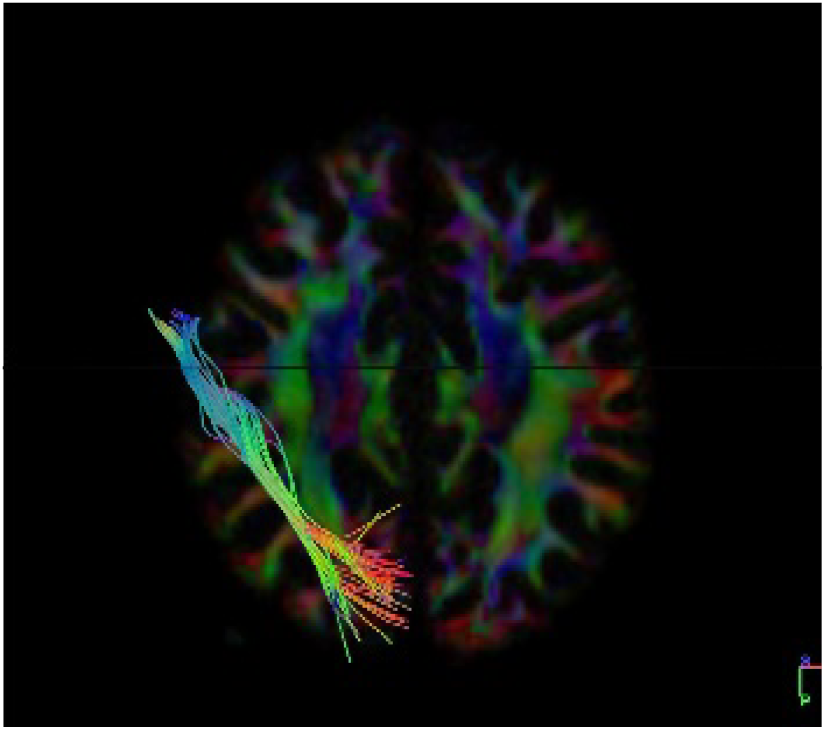
Axial Section Dataset subject – 1029 Male Right Side

**(Figure-2).**
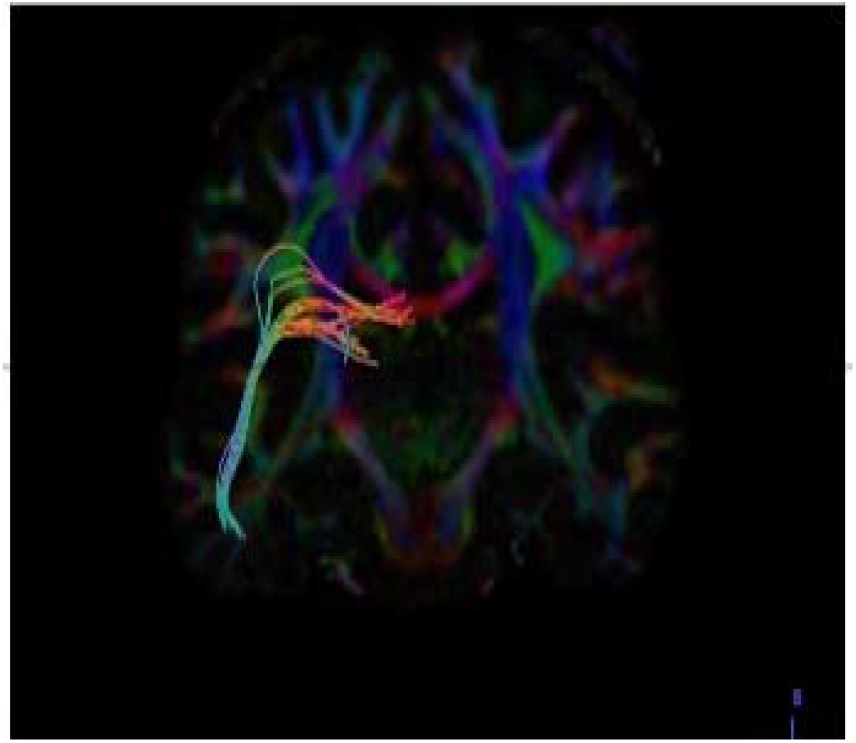
Coronal Section Dataset subject – 1029 Male Left Side

##### Graphical Representation

**(Graph-1)** plot in between number of tracts on x-axis and 16 subjects in y-axis to compare both left and right side of male datasets graphically. And we found that on an average right side is having a greater number of tracts when compared with left side in male subjects.

**Graph-1:**
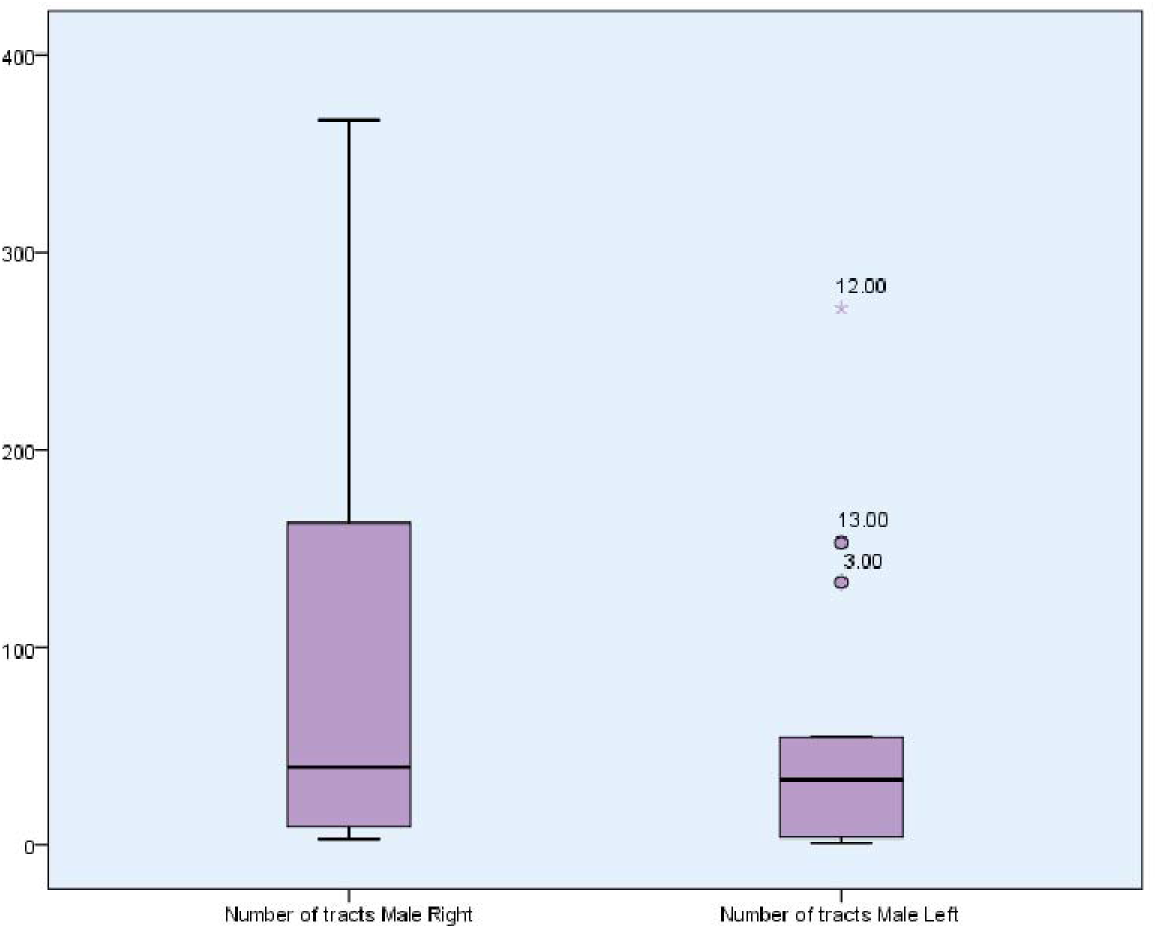
Comparison of Number of Tracts Right and left Ventral Stream Visual Pathway in Male Visual Pathway in Male. Normally in visual statistical learning when the right hemisphere is the dominant advantage of visual spatial integration and when the left hemisphere is also dominant, it is related to conceptual knowledge from patterns of covariation [92]. So, we conclude that there is no complete right side or complete left side dominance of ventral stream visual pathway in females, since if one side has less fibres the other side is balanced with its equalising fibres the same vice versa changes were seen if there are more fibres in one side, the other side is balanced with a less number of fibres.

#### b) Female subjects Right & Left

The Fibres were traced and observed for neural structural connectivity on both right and left side in all sixteen healthy adult female datasets **(Table 2).**

**Table 2:**
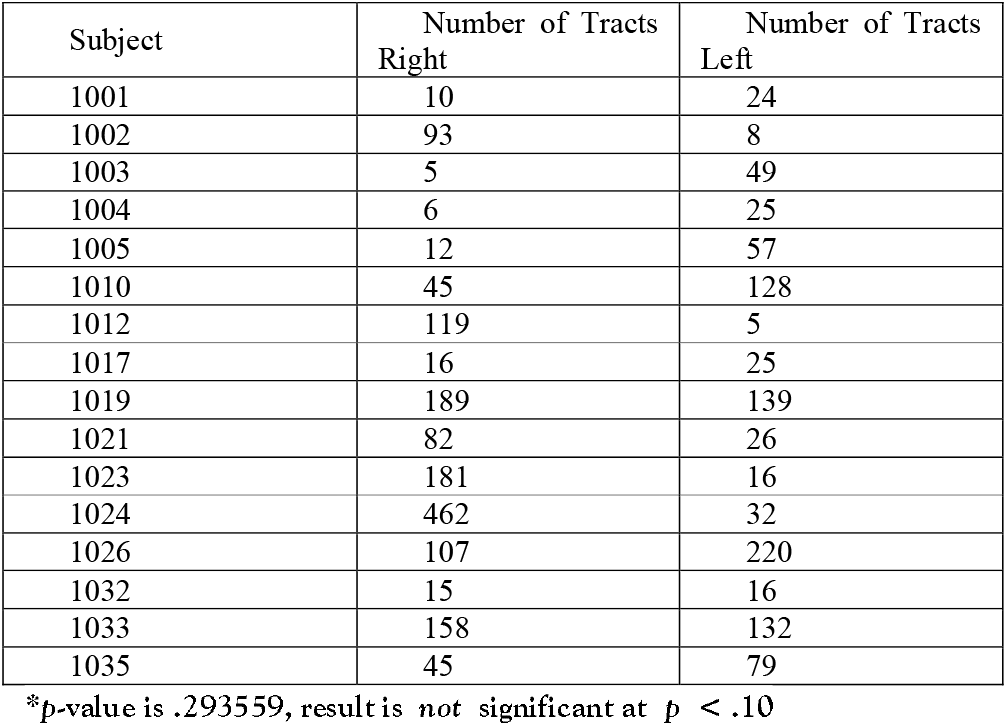
Number of Tracts Right and Left in Female *

| Subject | Number of Tracts<br>Right | Number of Tracts<br>Left |
| --- | --- | --- |
| 1001 | 10 | 24 |
| 1002 | 93 | 8 |
| 1003 | 5 | 49 |
| 1004 | 6 | 25 |
| 1005 | 12 | 57 |
| 1010 | 45 | 128 |
| 1012 | 119 | 5 |
| 1017 | 16 | 25 |
| 1019 | 189 | 139 |
| 1021 | 82 | 26 |
| 1023 | 181 | 16 |
| 1024 | 462 | 32 |
| 1026 | 107 | 220 |
| 1032 | 15 | 16 |
| 1033 | 158 | 132 |
| 1035 | 45 | 79 |

\*p-value is .293559, result is *not* significant at $p < .10$

Dataset numbers 1002, 1012, 1019, 1021, 1023, 1024, 1033 shows greater number of tracts at right side. Dataset numbers 1001, 1003, 1004, 1005, 1010, 1026, 1035 shows greater number of tracts on left side. Datasets numbers 1017 and 1032 does not show much difference between the tracts. Therefore, from the table given below, 6 datasets have more fibers on right and 6 datasets have more fibers on left and no difference was seen in 2 datasets but the variation in the count of fibers are more in right side **(dataset number 1021).**

When the results were statistically compared by performing independent t–test, no significant difference was seen between the number of tracts on right and left side of female subjects.

**(Figure-3).**
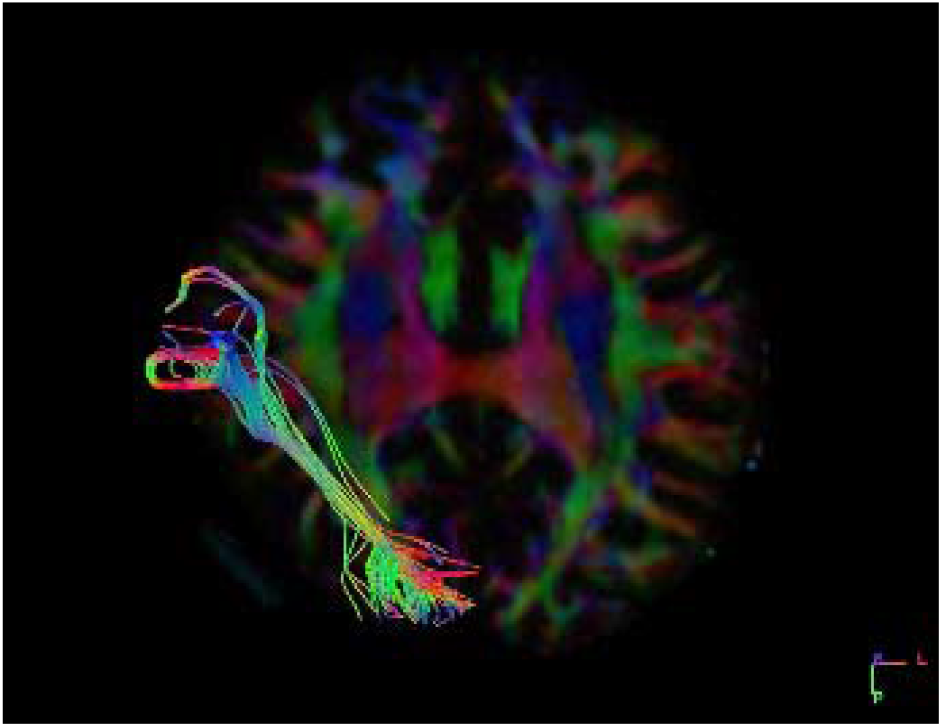
Axial Section Dataset subject - 1021 Female Right Side

**(Figure-4).**
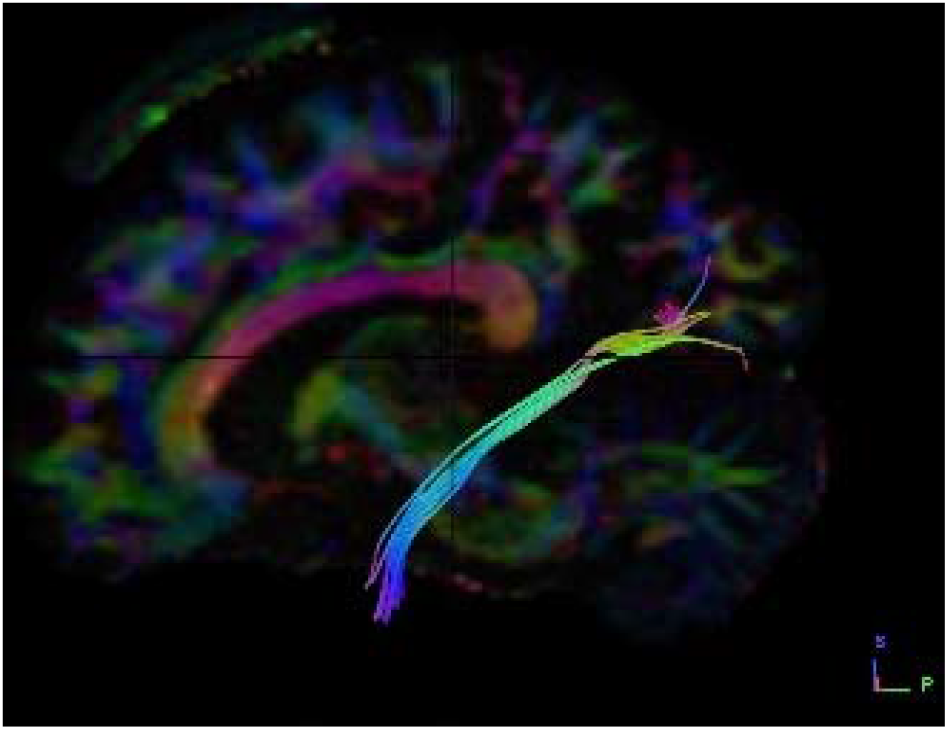
Sagittal Section Dataset subject - 1021 Female Left Side

##### Graphical Representation

**(Graph-2)** plot showing number of tracts on x-axis and 16 subjects on y-axis to compare both left and right side of female datasets graphically. We found that on average, the right side is having a greater number of tracts when compared with left side in female subjects.

**Graph-2:**
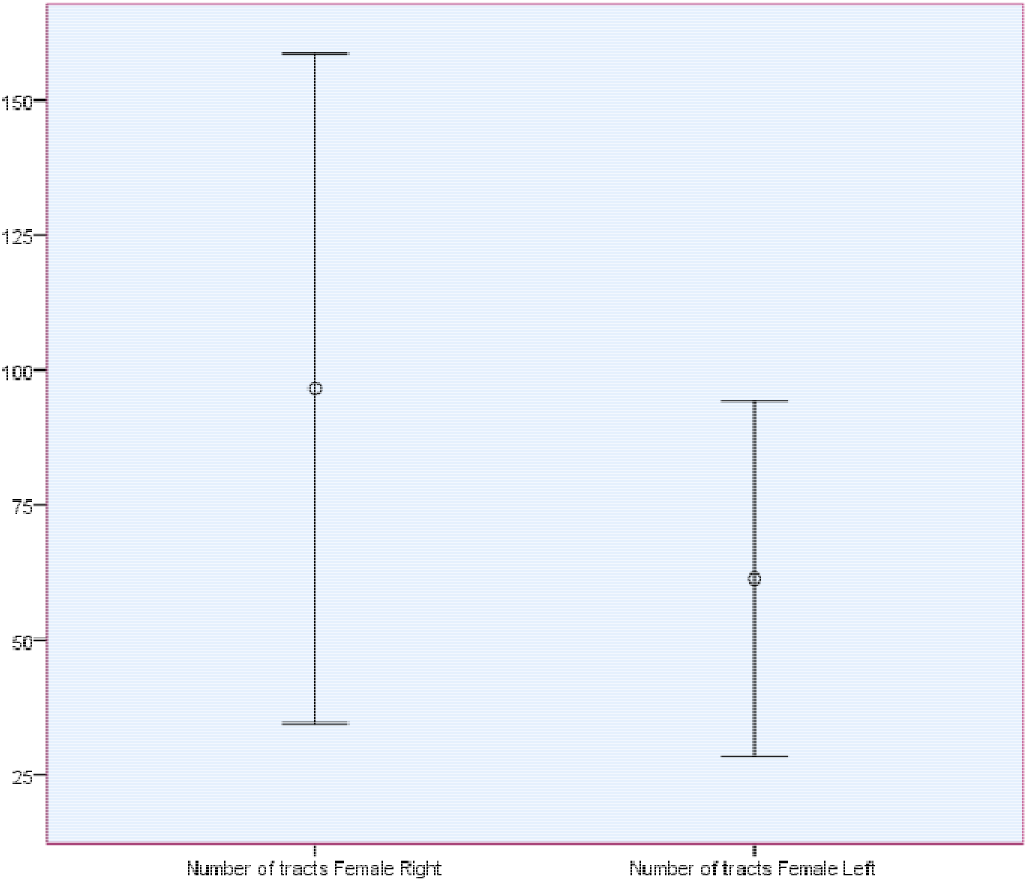
Comparison of Number of Tracts Right and left Ventral Stream Visual Pathway in Female. So, we conclude that there is no complete right side or complete left side dominance of ventral stream visual pathway in females since if one side has less fibres, the other side is balanced with its equalising fibres and same vice versa changes were seen if there are more fibres on one side, the other side is balanced with a smaller number of fibres.

#### C) Right Side Both Male and Female Subjects

The Fibres were traced and observed for neural structural connectivity on both right and left side in all sixteen healthy adult female datasets **(Table 3)**.

**Table 3:**
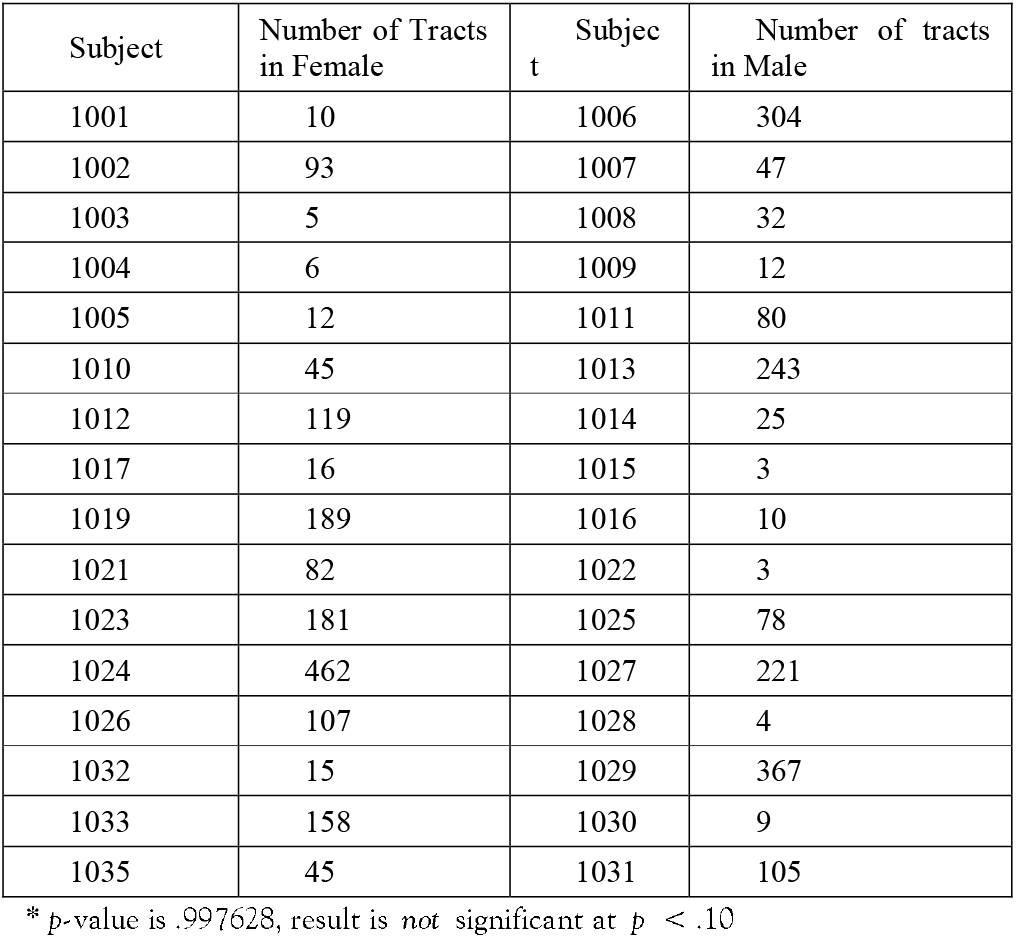
Number of Tracts Right Female and Male *.

When the results were statistically compared by performing independent t–test, no significant difference was seen between the number of tracts on right and left side of female subjects.

Dataset numbers 1002, 1010, 1012, 1019, 1023, 1024, 1026, 1033, 1035 shows greater number of tracts in females. Dataset numbers 1006, 1007, 1008, 1011, 1013, 1025, 1027, 1029, 1031 shows a greater number of tracts in males. Therefore, from the table given below, 9 datasets have more fibers in male and 9 datasets have more fibers in female but the variation in the count of fibers are more seen in right side of male **(dataset number 1024)**.

**(Figure-5).**
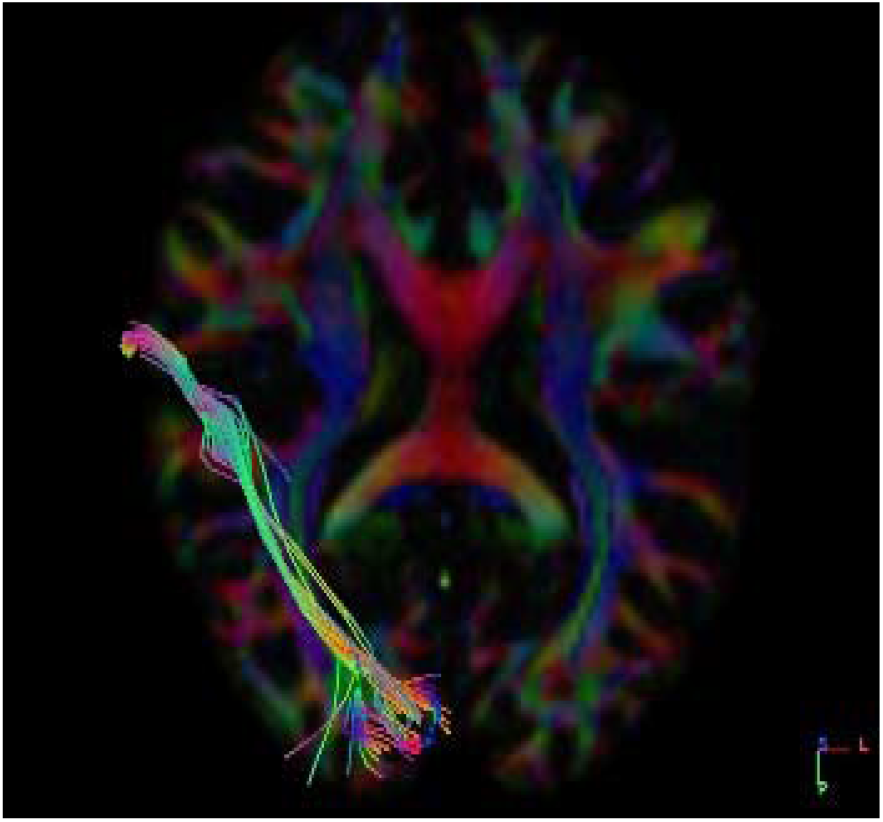
Axial Section Dataset subject - 1024 Male Right Side

**(Figure-6).**
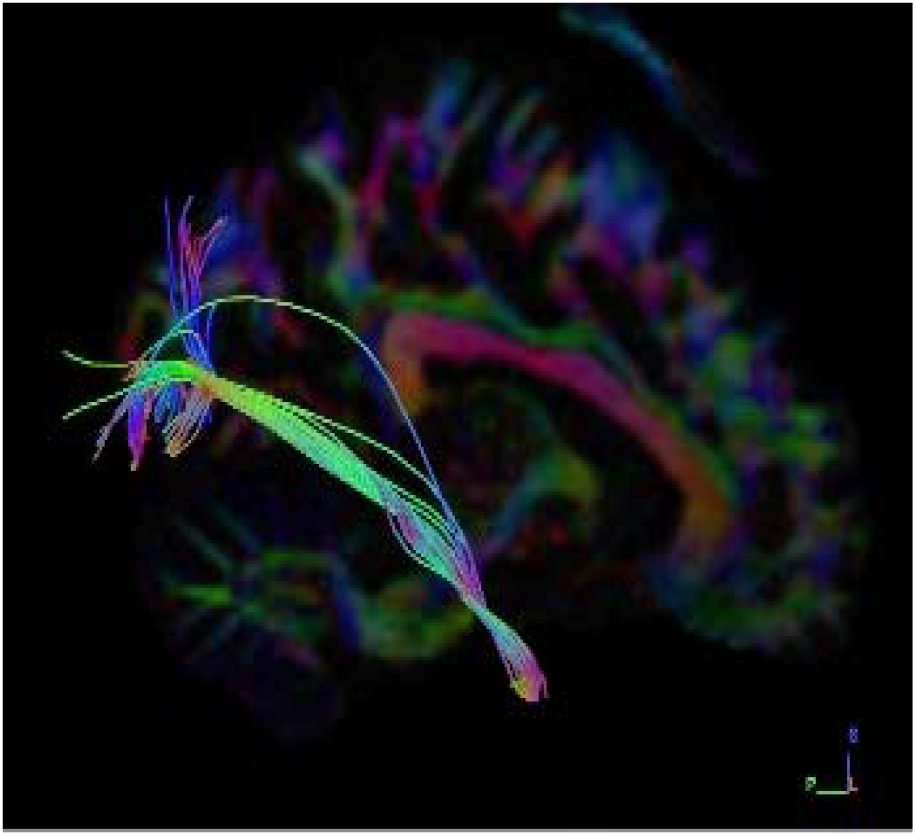
Sagittal Section Dataset subject – 1024 Female Right Side

##### Graphical Representation

**(Graph-3)** plot showing number of tracts on x-axis and **16** subjects on y-axis to compare right side of both male and female datasets graphically. Males typically outperform females on spatial tasks [93] and we found that on an average right side of males having a greater number of tracts when compared with left side in female subjects.

**Graph-3:**
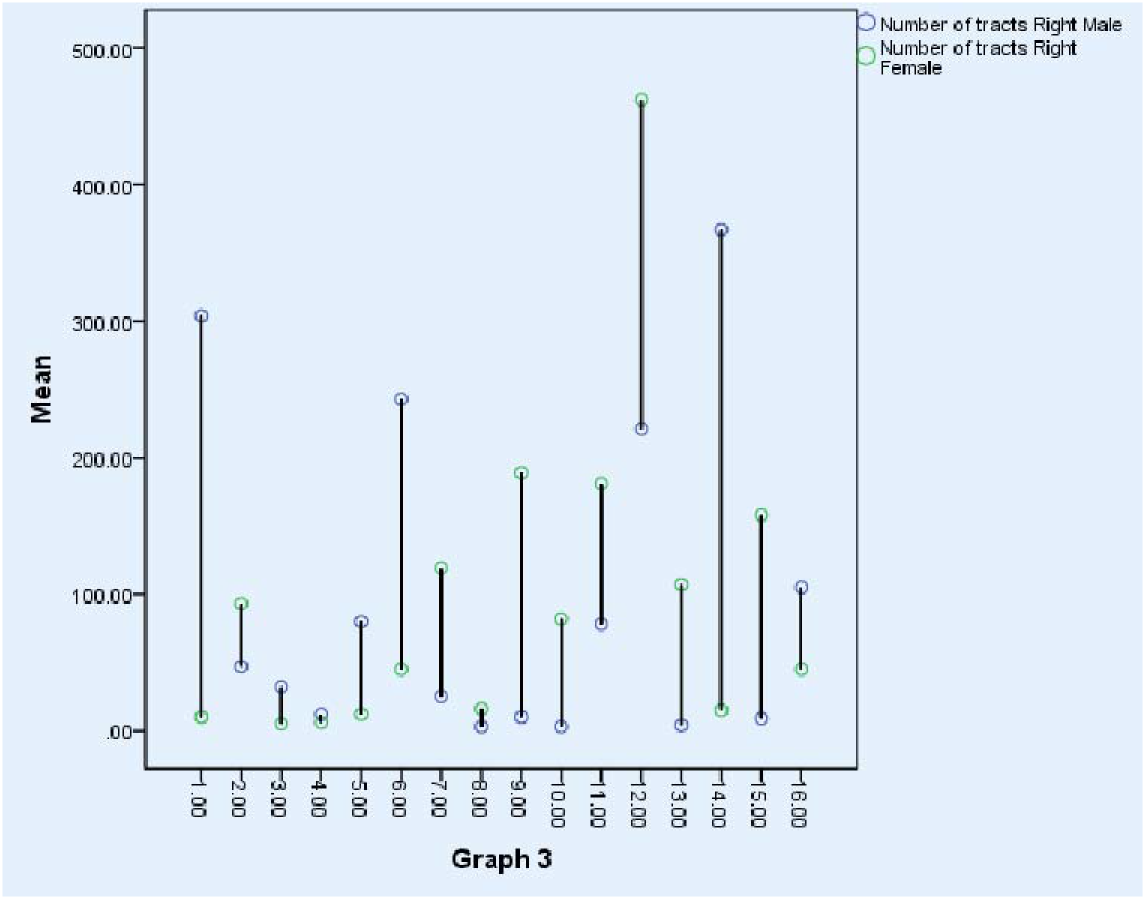
Comparison of Number of Tracts Right side Ventral Stream Visual Pathway in Female and Male. So, we conclude that there is no right side or left side dominance of ventral stream visual pathway in females, since if one side has less fibres, the other side is balanced with its equalising fibres on the other side and same vice versa changes were seen if there are more fibres in one side, the other side is balanced with less number of fibres.

#### d) Left Side Male and Female Subjects

The Fibres were traced and observed for neural structural connectivity on both right and left side in all sixteen healthy adult male datasets **(Table 4).** Dataset numbers 1003, 1005, 1010, 1019, 1026, 1033, 1035 shows greater number of tracts in female. Dataset numbers 1008, 1009, 1022, 1025,1027, 1028,1029, 1030, 1031 shows a greater number of tracts in male. Therefore, from the table given below, 9 datasets have more fibers at male and 7 datasets have more fibers in but the variation in the count of fibers are more on left side of male **(dataset number 1024).**

**Table 4:** Number of Tracts Female and Male Left *

| Subject | Number of Tracts Female | Subject | Number of tracts Male |
| --- | --- | --- | --- |
| 1001 | 24 | 1006 | 1 |
| 1002 | 8 | 1007 | 6 |
| 1003 | 49 | 1008 | 133 |
| 1004 | 25 | 1009 | 21 |
| 1005 | 57 | 1011 | 11 |
| 1010 | 128 | 1013 | 17 |
| 1012 | 5 | 1014 | 1 |
| 1017 | 25 | 1015 | 1 |
| 1019 | 139 | 1016 | 2 |
| 1021 | 26 | 1022 | 55 |
| 1023 | 16 | 1025 | 48 |
| 1024 | 32 | 1027 | 272 |
| 1026 | 220 | 1028 | 153 |
| 1032 | 16 | 1029 | 54 |
| 1033 | 132 | 1030 | 45 |
| 1035 | 79 | 1031 | 45 |

\*p-value is .765221, result is *not* significant at $p < .10$

**(Figure-7).**
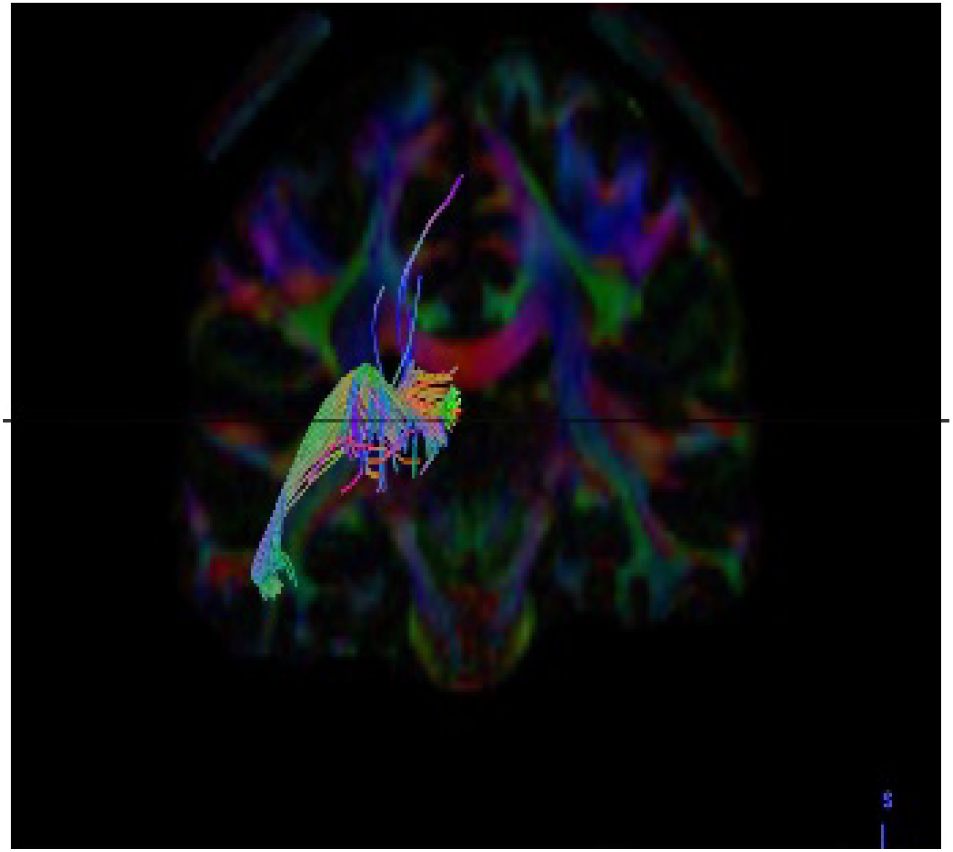
Coronal Section Dataset subject – 1024 Male Left Side

**(Figure-8).**
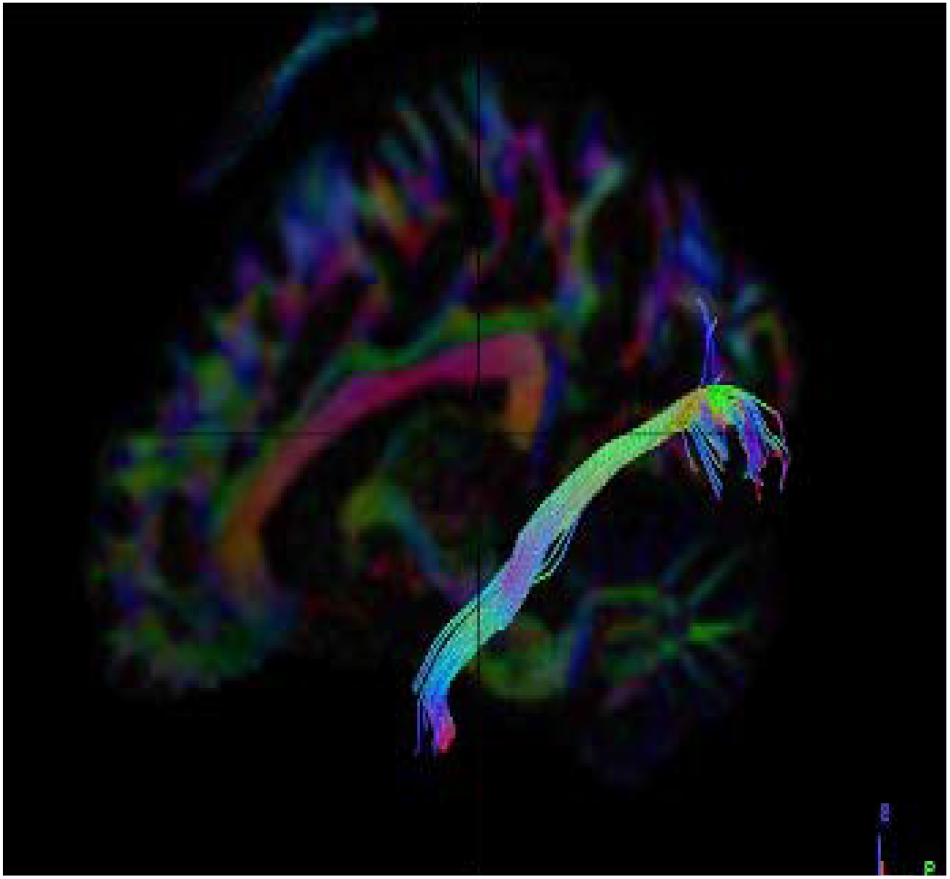
Sagittal Section Dataset subject – 1024 Female left Side

##### Graphical Representation

**(Graph-4)** plot showing number of tracts on x-axis and 16 subjects on y-axis to compare left side of both male and female datasets graphically. Males typically outperform females on spatial tasks [93] and we found that on an average left side is having a greater number of tracts when compared with female subjects.

**Graph-4:**
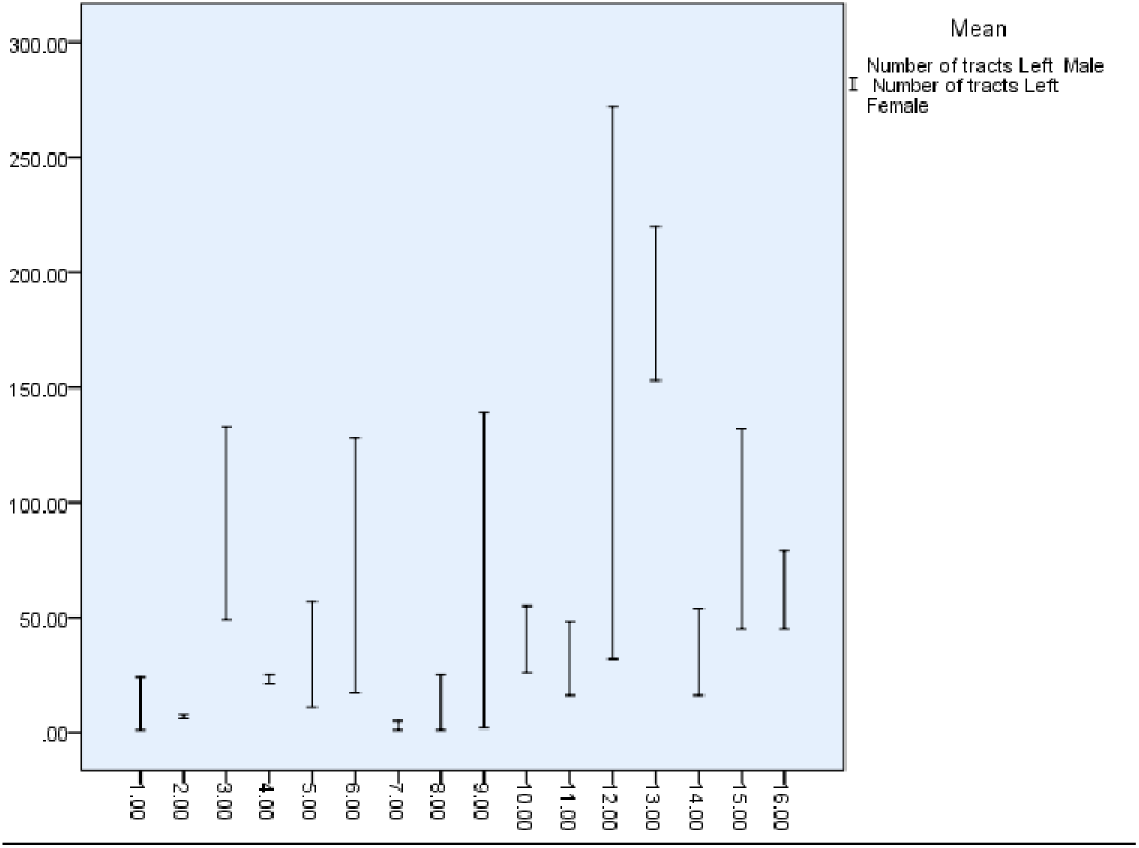
Comparison of Number of Tracts Left side Ventral Stream Visual Pathway in Female and Male. So, we conclude that there is no right side or left side dominance of ventral stream visual pathway in females, since if one side has less fibres, the other side is balanced with its equalising fibres on the other side and same vice versa changes were seen if there are more fibres in one side the other side is balanced with a smaller number of fibres.

### ii) Volume of Tracts

Tract volume is the total volume occupied by each fibre. Total volume is otherwise called the total number of voxels that were assigned as containing neural fibres. In correlation with our study, decrease or regression in the volume of the tracts might lead to clinical manifestations such as Visual Agnosia or Less efficiency of perceiving any object. The comparison of volume of Fibres is mostly done between the seed and the end region under four parameters;

a. Male Right and Left Side
b. Female Right and Left side
c. Right side Male and Female
d. Left side Male and Female

#### a) Male Subjects Right & Left

The Fibres were traced and observed for neural structural connectivity on both right and left side in all sixteen healthy adult male datasets **(Table 5).**

**Table 5:** Volume of Tracts male Right and Left*

| Subject | Volume of Tracts<br>Right | Volume of Tracts<br>Left |
| --- | --- | --- |
| 1006 | 1171.12 | 2838.37 |
| 1007 | 4920.75 | 1218.37 |
| 1008 | 1127.25 | 4941 |
| 1009 | 1150.87 | 1782 |
| 1011 | 1812.37 | 5629.5 |
| 1013 | 4714.87 | 7668 |
| 1014 | 6709.5 | 1090.13 |
| 1015 | 2234.25 | 2652.75 |
| 1016 | 5082.75 | 3526.88 |
| 1022 | 6024.38 | 2794.5 |
| 1025 | 8893.13 | 4131 |
| 1027 | 4154.63 | 5686.88 |
| 1028 | 1947.37 | 2737.12 |
| 1029 | 6901.87 | 7374.37 |
| 1030 | 4714.87 | 8721 |
| 1031 | 1171.12 | 2838.37 |

\*p-value is .293559, result is *not* significant at $p < .10$

Dataset number 1029 has the greatest volume right side and dataset number 1006 and 1031 has the least volume of tracts at right. Dataset number 1029 has greatest volume of tracts on left and dataset number 1014 has the least volume of tracts at left. Therefore, from the table given below, 7 datasets have more fibers at right and 9 datasets have more fibers at left but the variation in the count of fibers are more in left Side **(dataset number 1029).**

When the results were statistically compared by performing independent t–test, no significant difference was seen between the volume of tracts on right and left side of male subjects.

**(Figure-9).**
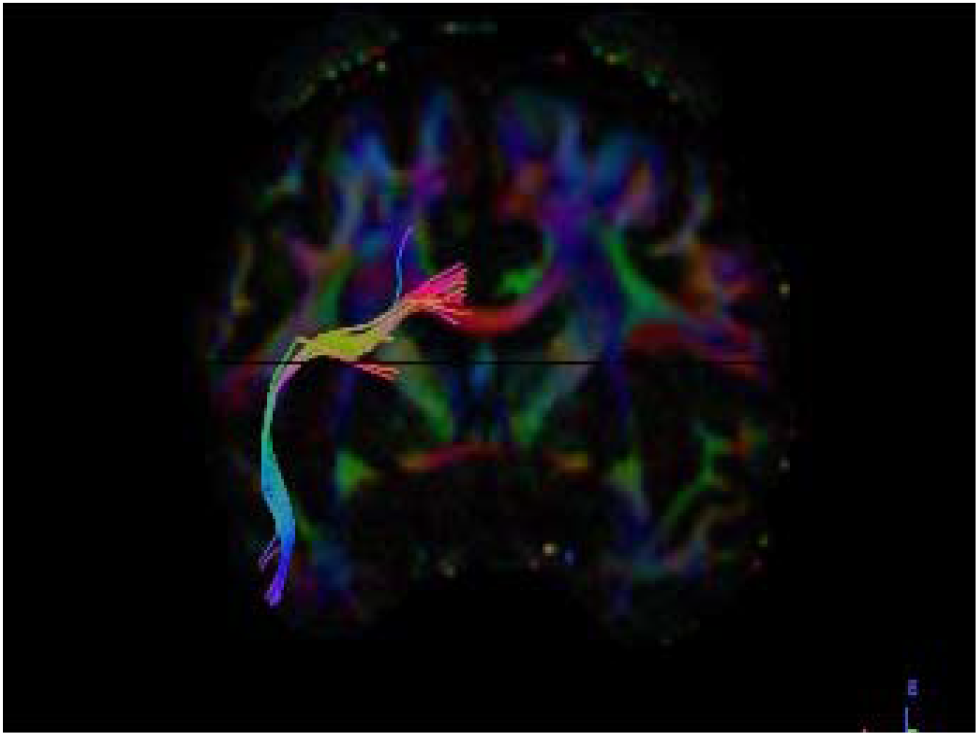
Coronal Section Dataset subject– 1029 Male Right Side

**(Figure-10).**
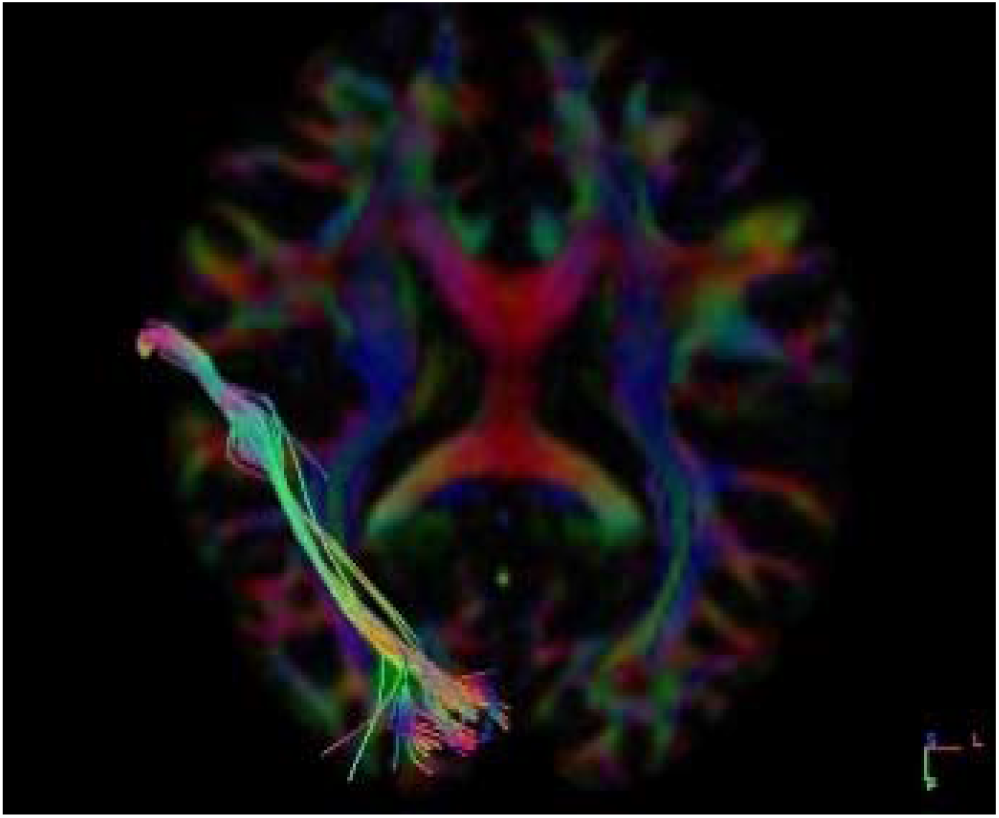
Coronal Section Dataset subject– 1029 Female Right Side

##### Graphical Representation

**(Graph-5)** plot showing number of tracts on x-axis and 16 subjects on y-axis to compare both left and right side of female datasets graphically. And we found that on an average left side is having a greater number of tracts when compared with right side in male subjects.

**Graph-5:**
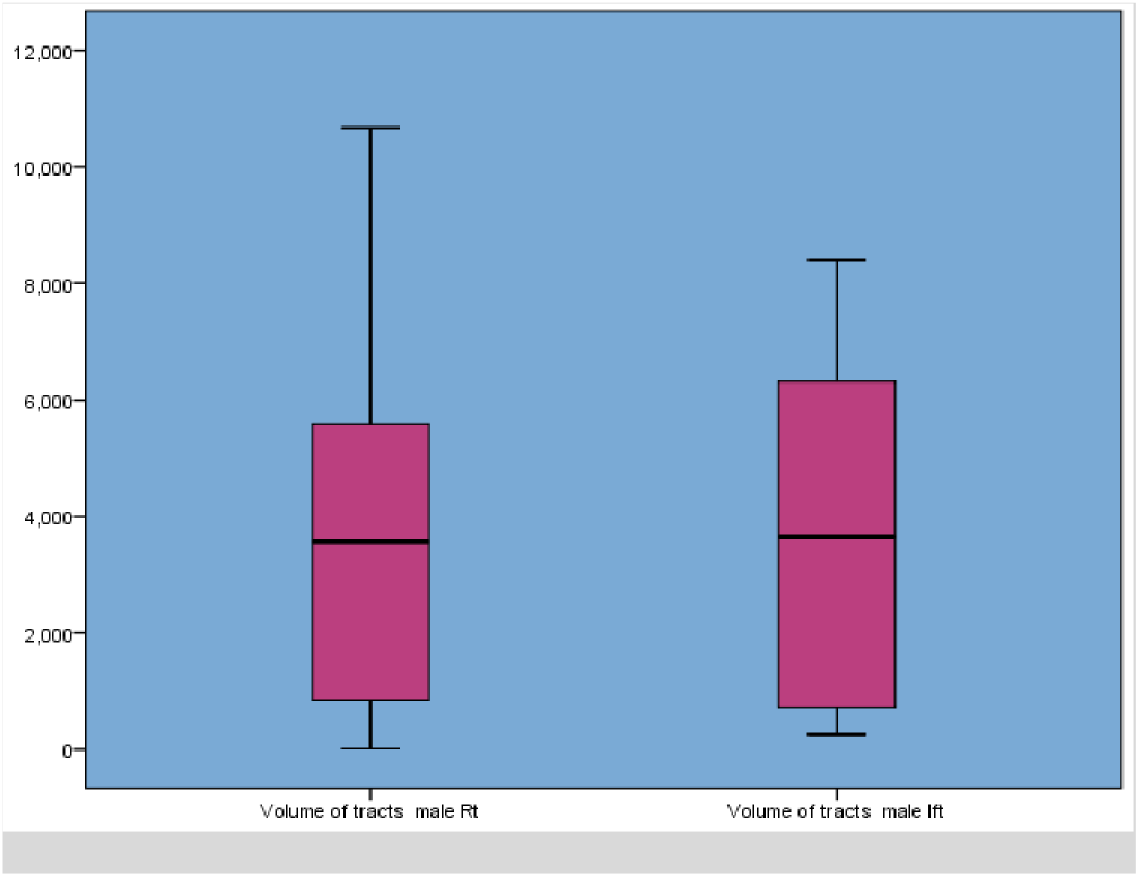
Comparison of Volume of Tracts Right & Left Side of Ventral Stream Visual Pathway in Female. We conclude that there is no complete right side or complete left side dominance of ventral stream visual pathway in females, since if one side has less fibres, the other side is balanced with its equalising fibres and same vice versa changes were seen if there are more fibres in one side the other side is balanced with less number of fibres.

#### b) Right and Left side of Female subjects

The Fibres were traced and observed for neural structural connectivity on both right and left side in all sixteen healthy adult female datasets **(Table 2).**

Dataset numbers 1002, 1003, 1005, 1010, 1012, 1021, 1023, 1024 show greater volume of tracts at right side. Dataset numbers 1004, 1005, 1010, 1021, 1024, 1026, 1033, 1035 show a greater number of tracts on left side. Datasets number 1010 shows more volume of fibers in right side and 1033 shows least volume of fibers on the right side. Dataset number 1026 shows more volume of fibers in the left side and 1001 shows least fibers in the left side. Therefore, the variation in the volume of fibers are more on left side of female subject **(dataset number 1021).**

**Table 6:**
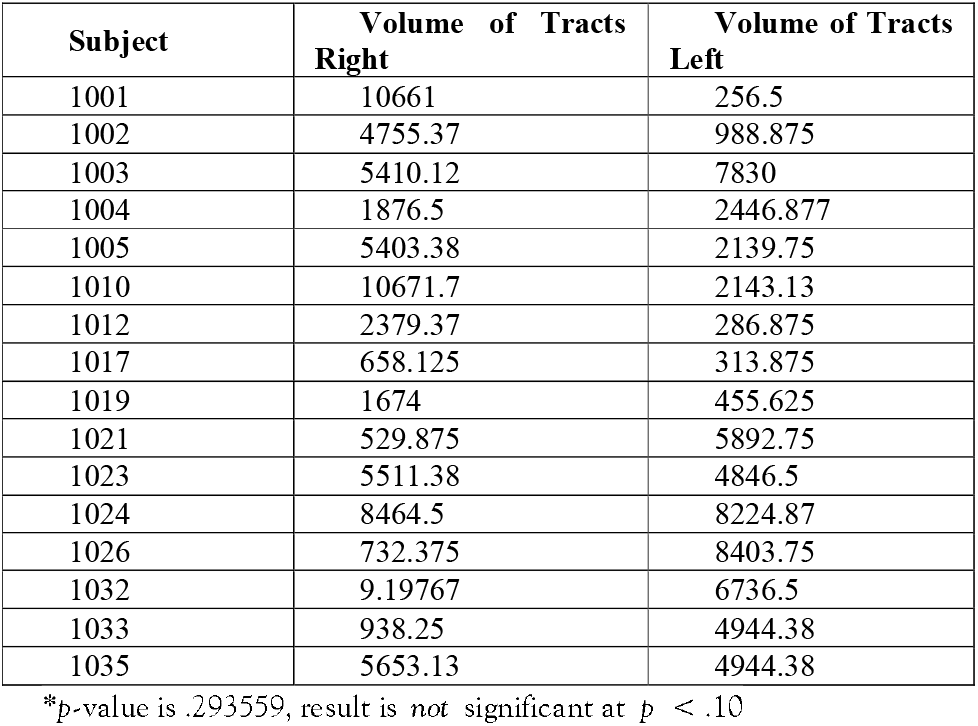
Volume Of tracts Female Right and Left *

When the results were statistically compared by performing independent t–test, no significant difference was seen between the volume of tracts in right and left side of female subjects.

**(Figure-11).**
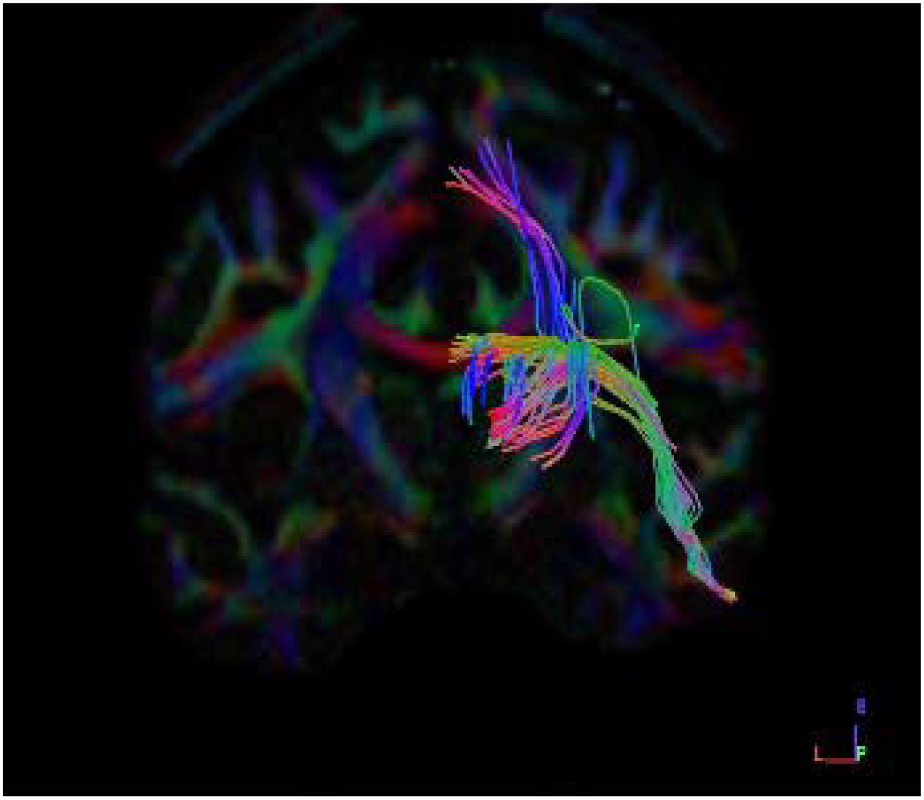
Coronal Section Dataset subject– 1021 Female Right Side

**(Figure-12).**
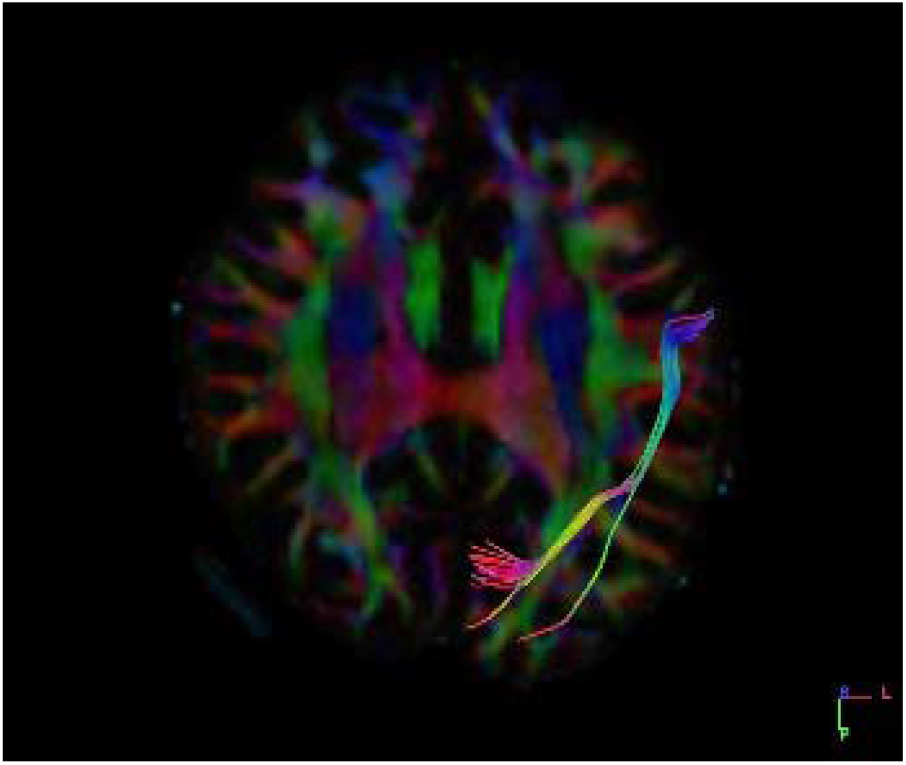
Coronal Section Dataset subject– 1021 Female Left Side

##### 1. Graphical Representation

**(Graph-6)** plot showing Volume of tracts on x-axis and 16 subjects on y-axis to compare both left and right side of female datasets graphically. We found that on average, the left side is having a greater number of tracts when compared with right side in female subjects.

**Graph-6:**
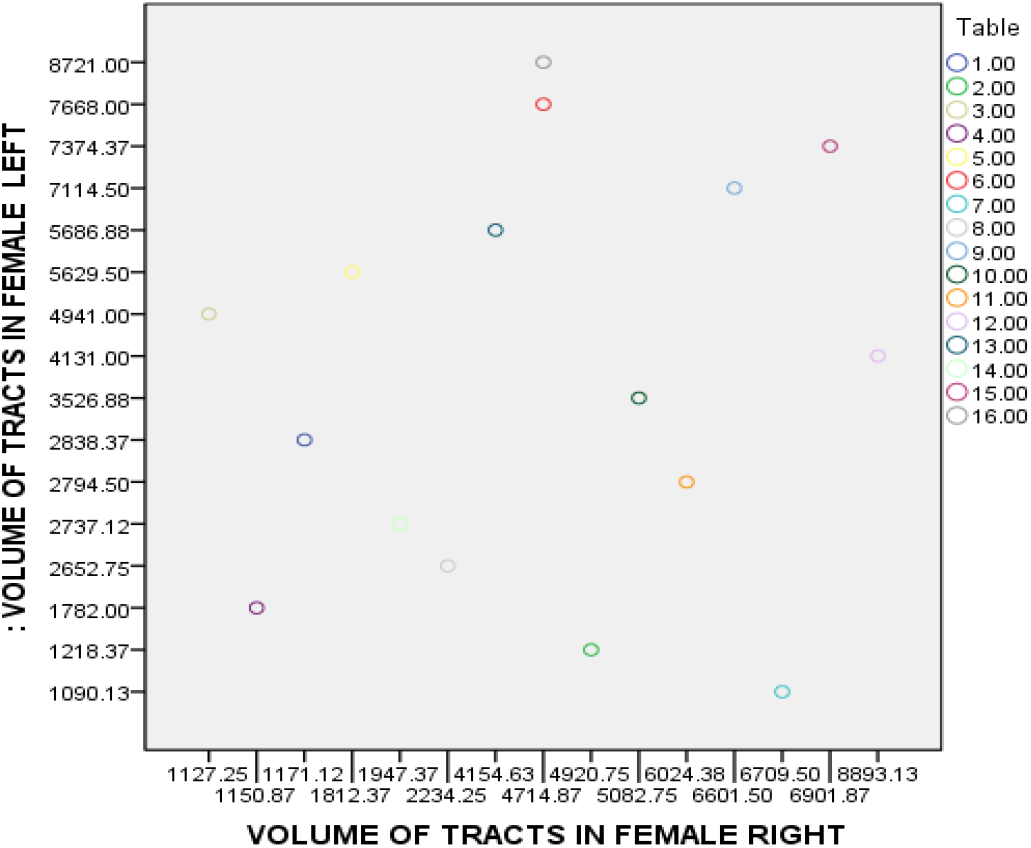
Comparison of Volume of Tracts Right & Left side of Ventral Stream Visual Pathway in Male. We conclude that there is no complete right side or complete left side dominance of ventral stream visual pathway in females, since if one side has less fibres the other side is balanced with its equalising fibres and same vice versa changes were seen if there are more fibres in one side, the other side is balanced with a smaller number of fibres.

#### C) Right Side Both Male and Female Subjects

The Fibres were traced and observed for neural structural connectivity on both right and left side in all sixteen healthy adult female datasets **(Table 7).**

**Table 7:** Volume of tracts in Female and Male Right *

| Subject | Volume of Tracts Female | Subject | Volume of tracts Male |
| --- | --- | --- | --- |
| 1001 | 1171.12 | 1006 | 10661 |
| 1002 | 4920.75 | 1007 | 4755.37 |
| 1003 | 1127.25 | 1008 | 5410.12 |
| 1004 | 1150.87 | 1009 | 1876.5 |
| 1005 | 1812.37 | 1011 | 5403.38 |
| 1010 | 4714.87 | 1013 | 10671.7 |
| 1012 | 6709.5 | 1014 | 2379.37 |
| 1017 | 2234.25 | 1015 | 658.125 |
| 1019 | 6601.5 | 1016 | 1674 |
| 1021 | 5082.75 | 1022 | 529.875 |
| 1023 | 6024.38 | 1025 | 5511.38 |
| 1024 | 8893.13 | 1027 | 8464.5 |
| 1026 | 4154.63 | 1028 | 732.375 |
| 1032 | 1947.37 | 1029 | 9.19767 |
| 1033 | 6901.87 | 1030 | 938.25 |
| 1035 | 4714.87 | 1031 | 5653.13 |

\*p-value is .870655, result is *not* significant at $p < .10$

Dataset numbers 1002, 1010, 1012, 1019, 1023, 1024, 1026, 1033, 1035 shows greater volume of tracts on female. Dataset numbers 1006, 1007, 1008, 1011, 1013, 1025, 1027, 1029, 1031 shows a greater number of tracts on male. Therefore, from the table given below, 9 datasets have more fibers in male and 9 datasets have more fibers in female but the variation in the count of fibers are more seen in right Side of male **(dataset number 1024).**

When the results were statistically compared by performing independent t–test, no significant difference were seen between the volume of tracts in right side of male and female subjects.

**(Fig 13).**
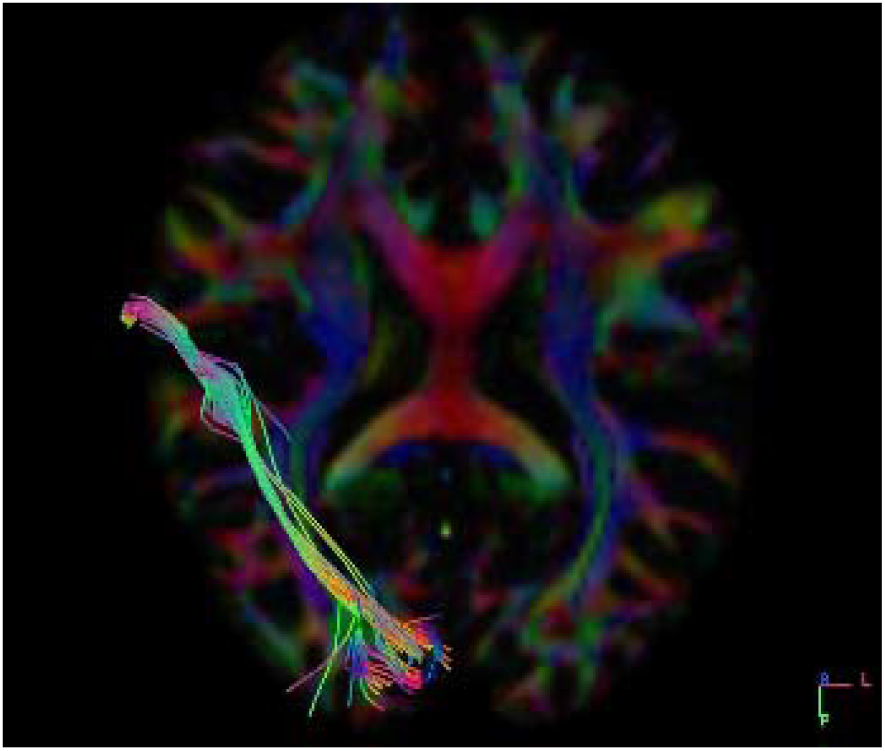
Axial Section Dataset subject – 1024 Female right Side

**(Fig 14).**
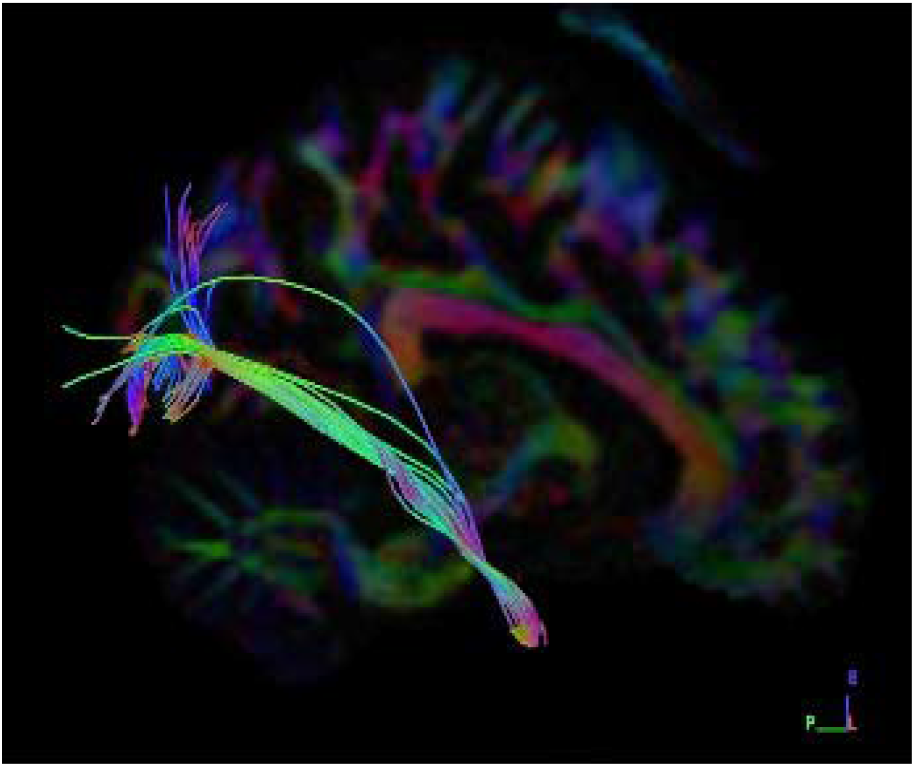
Sagittal section Dataset subject - 1024 Male ight side

##### Graphical Representation

**Graph-7:** plot showing Volume of tracts on x-axis and 16 subjects on y-axis to compare both left and right side of female datasets graphically. We found that on average, the right side is having a greater number of tracts when compared with right-side of both male and female subjects.

**Graph-7:**
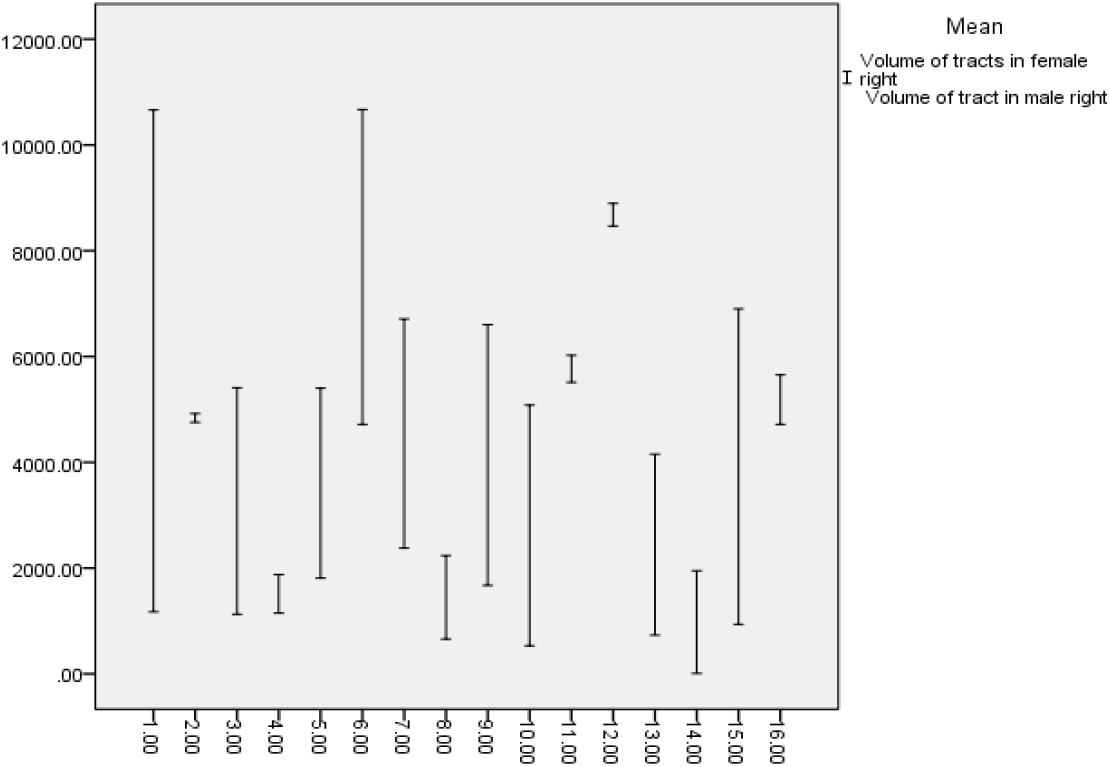
Comparison of Volume of Tracts in Right side of female and male Ventral Stream Visual Pathway. We conclude that no complete right-side dominance of ventral stream visual pathway is seen in male and female subjects, since if one side has less fibres the other side is balanced with its equalising fibres.

#### d) Left Side Male and Female Subjects

The Fibres were traced and observed for neural structural connectivity on both right and left side in all sixteen healthy adult male datasets **(Table-8).**

**Table 8:** Volume of tracts in Female and Male Left *

| Subject | Volume of Tracts Female | Subject | Volume of tracts Male |
| --- | --- | --- | --- |
| 1001 | 2838.37 | 1006 | 256.5 |
| 1002 | 1218.37 | 1007 | 988.875 |
| 1003 | 4941 | 1008 | 7830 |
| 1004 | 1782 | 1009 | 2446.877 |
| 1005 | 5629.5 | 1011 | 2139.75 |
| 1010 | 7668 | 1013 | 2143.13 |
| 1012 | 1090.13 | 1014 | 286.875 |
| 1017 | 2652.75 | 1015 | 313.875 |
| 1019 | 7114.5 | 1016 | 455.625 |
| 1021 | 3526.88 | 1022 | 5892.75 |
| 1023 | 2794.5 | 1025 | 4846.5 |
| 1024 | 4131 | 1027 | 8224.87 |
| 1026 | 5686.88 | 1028 | 8403.75 |
| 1032 | 2737.12 | 1029 | 6736.5 |
| 1033 | 7374.37 | 1030 | 4944.38 |
| 1035 | 8721 | 1031 | 4944.38 |

\*p-value is .564097, result is not significant at $p < .10$

Dataset numbers 1003, 1005, 1010, 1019, 1026, 1033, 1035 shows greater Volume of tracts in females. Dataset numbers 1008, 1009, 1022, 1025, 1027, 1028, 1029, 1030, 1031 shows a greater number of tracts in male. Therefore, from the table given below, 9 datasets have more fibers at male and 7 datasets have more fibers but the variation in the count of fibers are more in left side of male **(dataset number 1024).**

When the results were statistically compared by performing independent t–test no significant difference was seen between the volume of tracts in left side of male and female subjects.

**(Fig 16).**
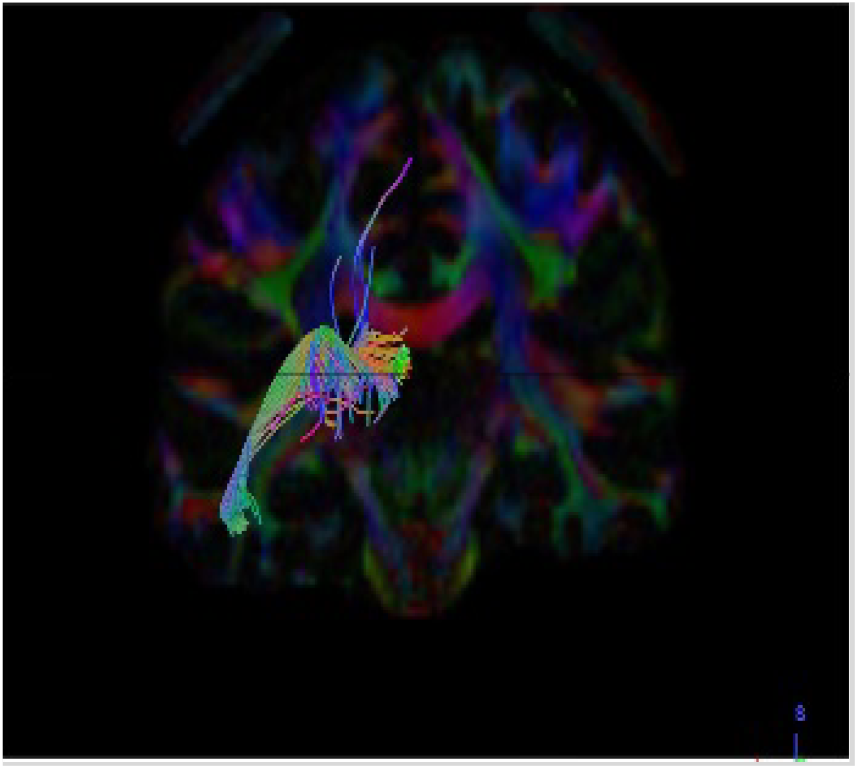
Coronal section Dataset Subject - 1028 Male left side

**(Fig 17).**
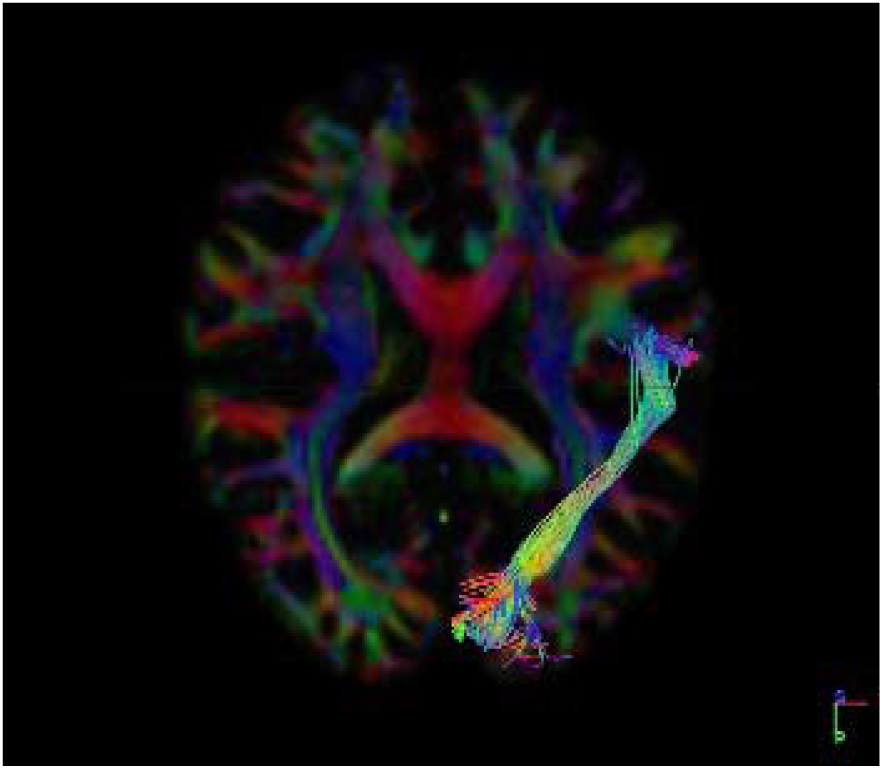
Axial Section Dataset Subject – 1026 Female left side

##### Graphical Representation

**(Graph-8)** plot showing Volume of tracts on x-axis and 16 subjects in y-axis to compare both left and right side of female datasets graphically. We found that on average, the left side of male is greater when compared with left -side of female subjects.

**Graph-8:**
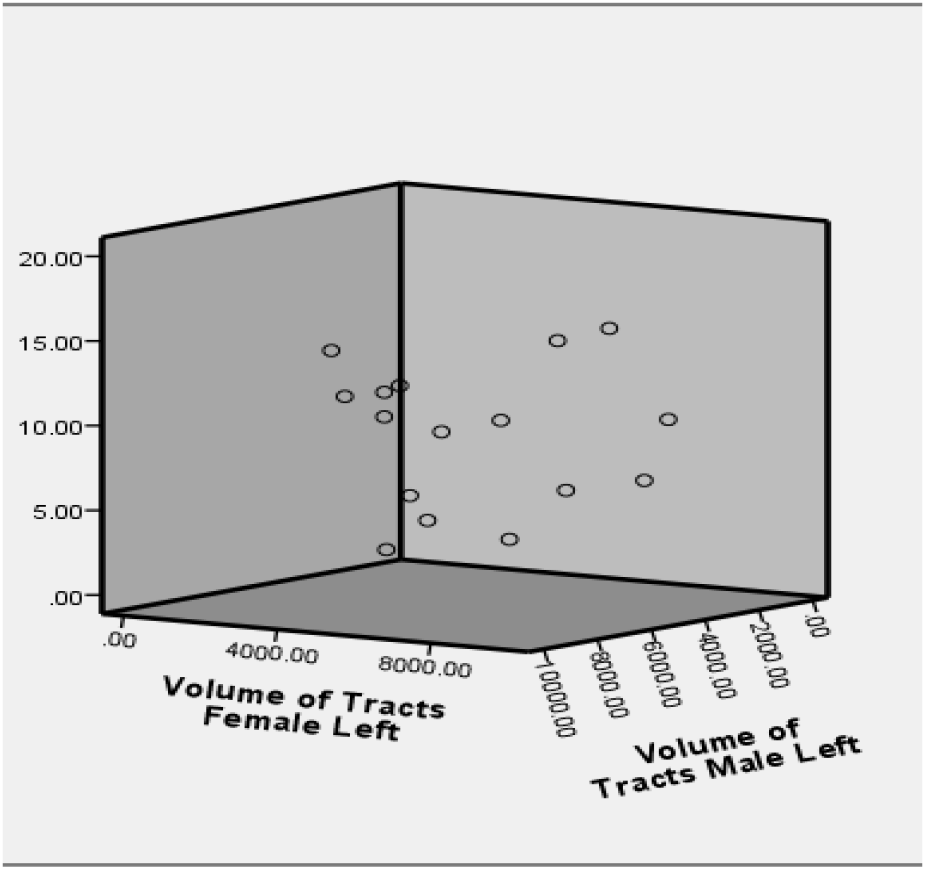
Comparison of Volume of Tracts in left side of female and male Ventral Stream Visual Pathway. We conclude that no complete left-side dominance of ventral stream visual pathway is seen in male and female subjects since if one side has less fibres the other side is balanced with its equalising fibres.

### iii) Mean Length of tracts

Tract mean length (mm) the average absolute deviation of a data from a central point. We are analysing tract mean length (mm) present in both the sexes (i.e. i) Male subjects and ii) Female subjects. The comparison of Mean Length of Fibres is mostly done between the seed and the end region under four parameters;

a. Male Right and Left Side
b. Female Right and Left side
c. Right side Male and Female
d. Left side Male and Female

#### a) Right and Left side of male subjects

The Fibres were traced and observed for neural structural connectivity on both right and left side in all sixteen healthy adult male datasets **(Table-9).**

**Table 9:** Mean Length of Fibres in Male Right and Left*

| **Subject** | ***Length of Tracts***<br>***Mean(mm) Right*** | ***Length of***<br>***Tracts Mean(mm)***<br>***Left*** |
| --- | --- | --- |
| 1006 | 112.114 | 86.2999 |
| 1007 | 91.6493 | 102.975 |
| 1008 | 111.561 | 97.361 |
| 1009 | 95.4516 | 91.4074 |
| 1011 | 94.2561 | 108.222 |
| 1013 | 99.3921 | 103.787 |
| 1014 | 102.589 | 119.844 |
| 1015 | 94.4761 | 97.666 |
| 1016 | 97.7039 | 88.922 |
| 1022 | 95.6268 | 113.468 |
| 1025 | 96.9707 | 122.641 |
| 1027 | 99.1497 | 119.758 |
| 1028 | 101.312 | 100.107 |
| 1029 | 101.808 | 109.768 |
| 1030 | 114.347 | 105.942 |
| 1031 | 107.152 | 105.942 |

\*p-value is .600727, result is *not* significant at $p < .10$

Dataset numbers 1006, 1007. 1011, 1013, 1014, 1016, 1027, 1029, 1031 shows greater number of tracts at right side. Dataset numbers 1008, 1009, 1015, 1022, 1027, 1028, 1030 shows a greater number of tracts on left side. Therefore, from the table given below, 9 datasets have more fibers at right and 7 datasets have more fibers at left but the variation in the mean length of fibers are more on right side **(dataset number 1030).**

When the results were statistically compared by performing independent t–test, no significant difference was seen between the mean length of tracts in right side of male subjects.

**(Fig 18).**
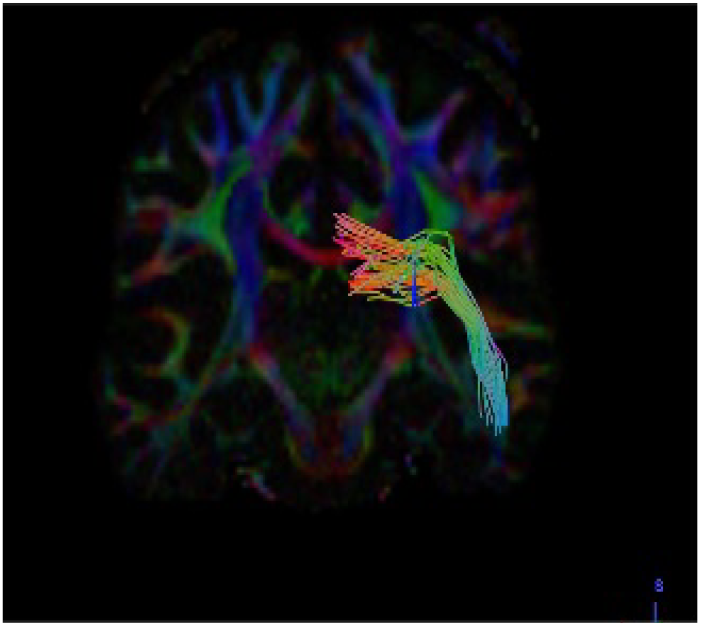
Coronal section Dataset Subject - 1030 Male right side

**(Fig 19).**
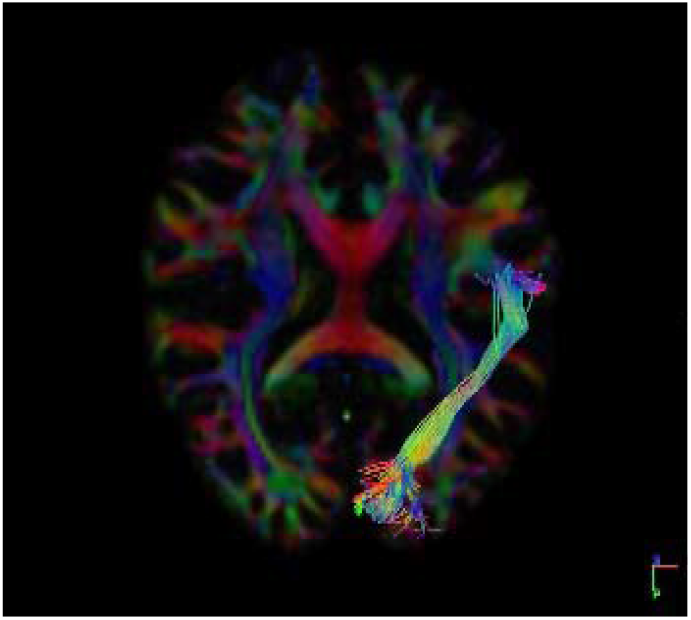
Axial section Dataset Subject - 1030 Male left side

##### Graphical Representation

**(Graph-9)** plot showing Mean Length of tracts on x-axis and 16 subjects on y-axis to compare both left and right side of female datasets graphically. And we found that on an average right side of male is greater when compared with left -side of male subjects.

**Graph-10:**
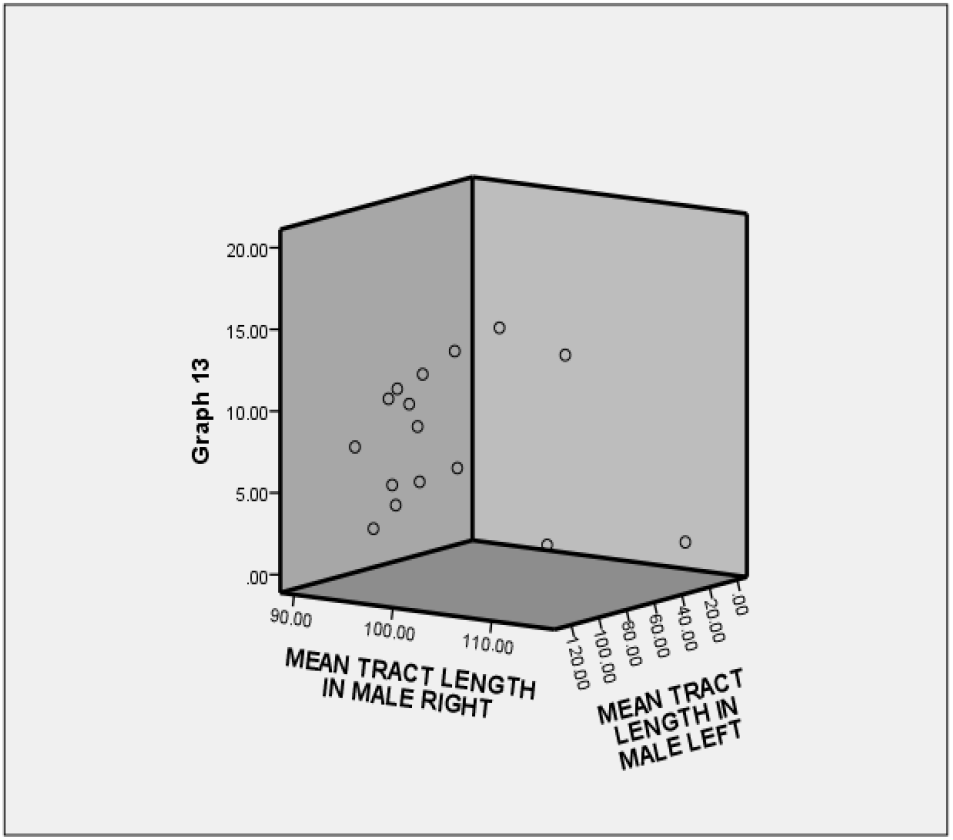
Comparison of Mean Length of Tracts in male right and left side. We conclude that there is no complete right-side dominance or complete left-side dominance of ventral stream visual pathway in male and female subjects, and since if one side has less fibres, the other side is balanced with its equalising fibres.

#### b) Right and Left side of Female subjects

The Fibres were traced and observed for neural structural connectivity on both right and left side in all sixteen healthy adult female datasets **(Table 10).** Dataset numbers 1002, 1012, 1019, 1021, 1023, 1024, 1033 shows greater number of tracts at right side. Dataset number 1001, 1003, 1004, 1005, 1010, 1026, 1035 shows a greater Mean length of tracts on left side. Datasets number 1017 and 1032 does not show any great difference between the tracts. Therefore, from the table given below, 6 datasets have more fibers at right and 6 datasets have more fibers at left and no difference has been seen in 2 datasets but the variation in the mean length of fibers are more in Left Side **(dataset number 1021).**

**Table 10:** Mean Length of Tracts in Female Right and Left *

| **Subject** | *Length of Tracts*<br>*Mean(mm) Right* | *Length of Tracts*<br>*Mean(mm) Left* |
| --- | --- | --- |
| 1001 | **106.752** | **94.8221** |
| 1002 | 109.728 | 103.318 |
| 1003 | 95.1141 | 95.034 |
| 1004 | 100.122 | 101.919 |
| 1005 | 93.776 | 101.059 |
| 1010 | 92.1292 | 94.5857 |
| 1012 | 99.1144 | 98.1627 |
| 1017 | 105.575 | 91.3946 |
| 1019 | 105.151 | 98.9259 |
| 1021 | 111.763 | 91.7771 |
| 1023 | 107.889 | 104.036 |
| 1024 | 107.667 | 98.6967 |
| 1026 | 84.4058 | 100.648 |
| 1032 | 105.667 | 114.255 |
| 1033 | 108.95 | 104.669 |
| 1035 | 92.1292 | 111.022 |

\*p-value is .600727, result is *not* significant at $p < .10$

When the results were statistically compared by performing independent t–test, no significant difference was seen between the number of tracts in right and left side of female subjects.

**(Fig 20).**
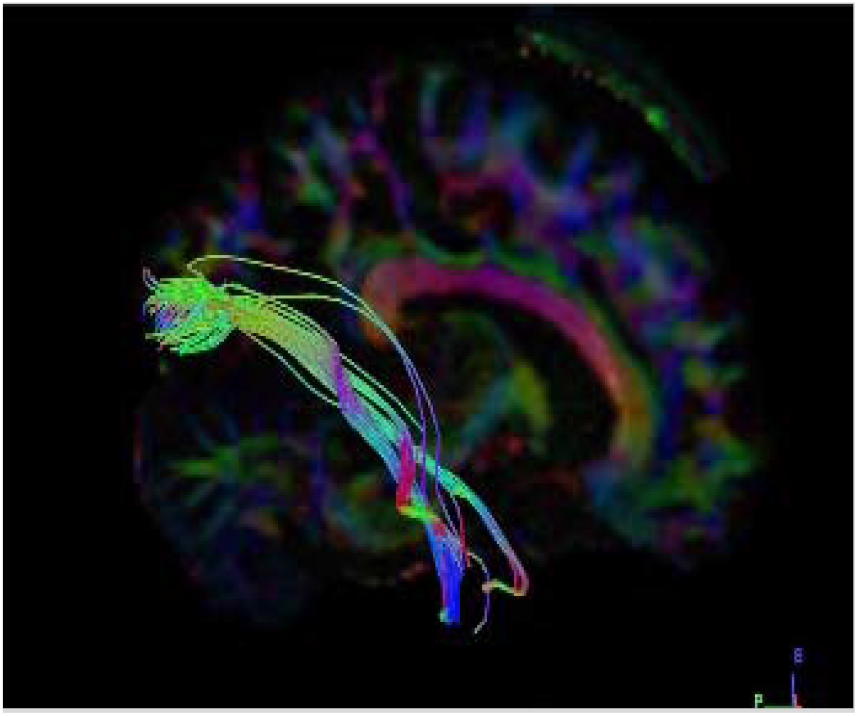
Sagittal section Dataset Subject - 1021 Female right side

**(Fig 21).**
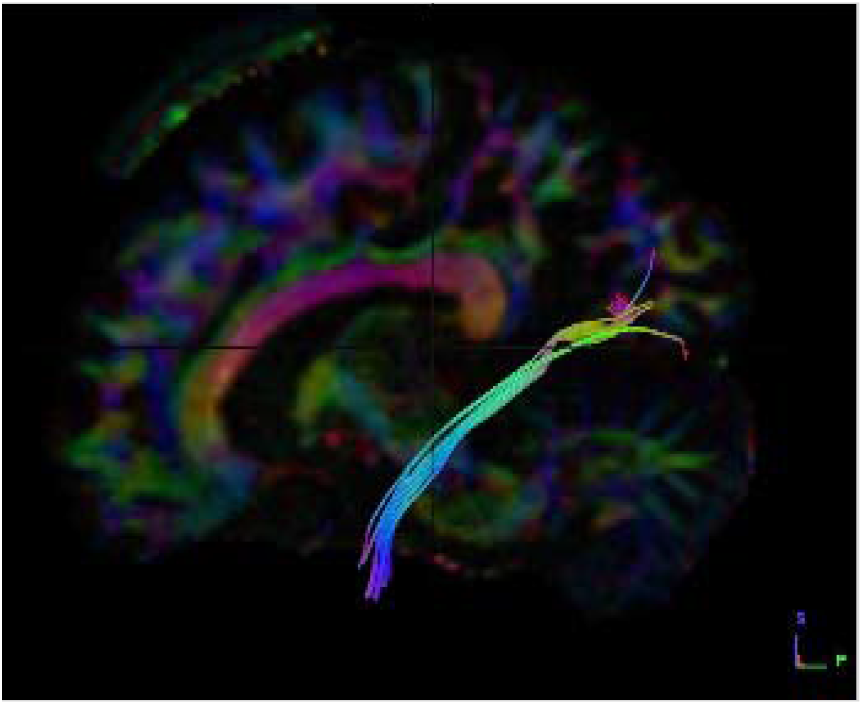
Sagittal section Dataset Subject - 1021 Female left side

##### Graphical Representation

**(Graph-10)** plot showing Mean Length of tracts on x-axis and 16 subjects on y-axis to compare both left and right side of female datasets graphically. And we found that on average, left side is having a greater number of tracts when compared with right side in female subjects.

**Graph-10:**
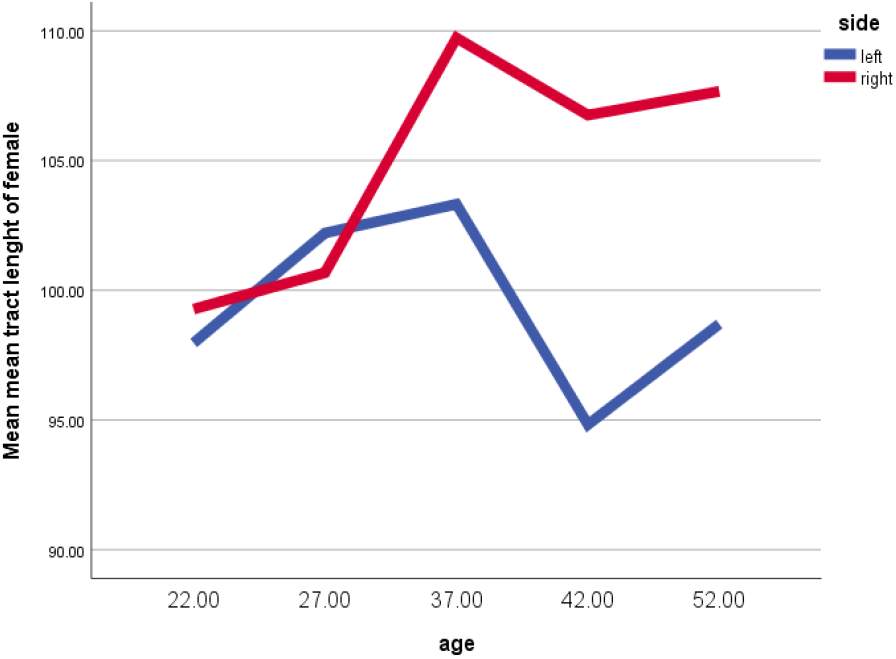
Comparison of Mean Length of Tracts on female right and left side. We conclude that there is no complete right side or complete left side dominance of ventral stream visual pathway in females. Hence, if one side has less fibres, the other side is balanced with its equalising fibres. Similar vice versa changes were seen if there are more fibres on one side, the other side is balanced with a smaller number of fibres.

#### c) Right side of Female and male subjects

The Fibres were traced and observed for neural structural connectivity on both right and left sides in all sixteen healthy adult female datasets **(Table-11)**.

**Table 11:** Mean length of fibres in Female and Male Right*

| **Subject** | *Length of Tracts*<br>*Mean(mm) Female* | *Length of Tracts*<br>*Mean(mm) Male* |
| --- | --- | --- |
| 1006 | **106.752** | 112.114 |
| 1007 | 109.728 | 91.6493 |
| 1008 | 95.1141 | 111.561 |
| 1009 | 100.122 | 95.4516 |
| 1011 | 93.776 | 94.2561 |
| 1013 | 92.1292 | 99.3921 |
| 1014 | 99.1144 | 102.589 |
| 1015 | 105.575 | 94.4761 |
| 1016 | 105.151 | 97.7039 |
| 1022 | 111.763 | 95.6268 |
| 1025 | 107.889 | 96.9707 |
| 1027 | 107.667 | 99.1497 |
| 1028 | 84.4058 | 101.312 |
| 1029 | 105.667 | 101.808 |
| 1030 | 108.95 | 114.347 |
| 1031 | 92.1292 | 107.152 |

\*The $p$ -value is .441763. The result is not significant at $p < .10$ .

Dataset numbers 1002, 1010, 1012, 1019, 1023, 1024, 1026, 1033, 1035 show greater number of tracts on female. Dataset numbers 1006, 1007, 1008, 1011, 1013, 1025, 1027, 1029, 1031 show a greater Mean Length of tracts in males. Therefore, from the table given below, 9 datasets have more fibers in male and 9 datasets have more fibers in female but the variation in the Mean Length of fibers are more seen in right Side of male subject **(dataset number 1024).**

When the results were statistically compared by performing independent t–test, no significant difference was seen between the Mean length of tracts in female and male subjects on right side.

**(Fig 20).**
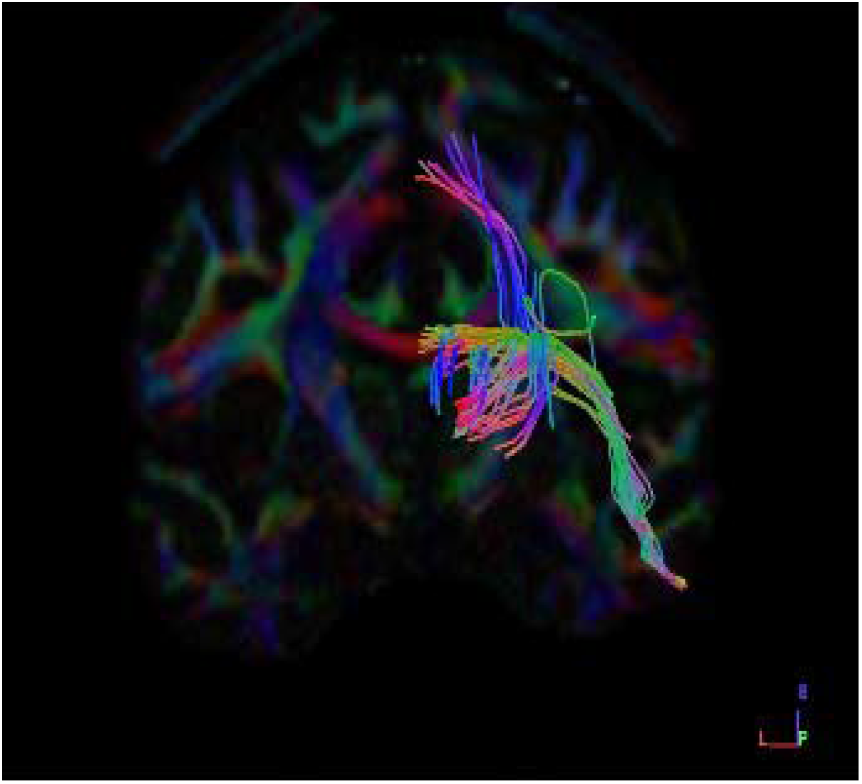
Coronal section Dataset Subject - 1024 Female right side

**(Fig 21).**
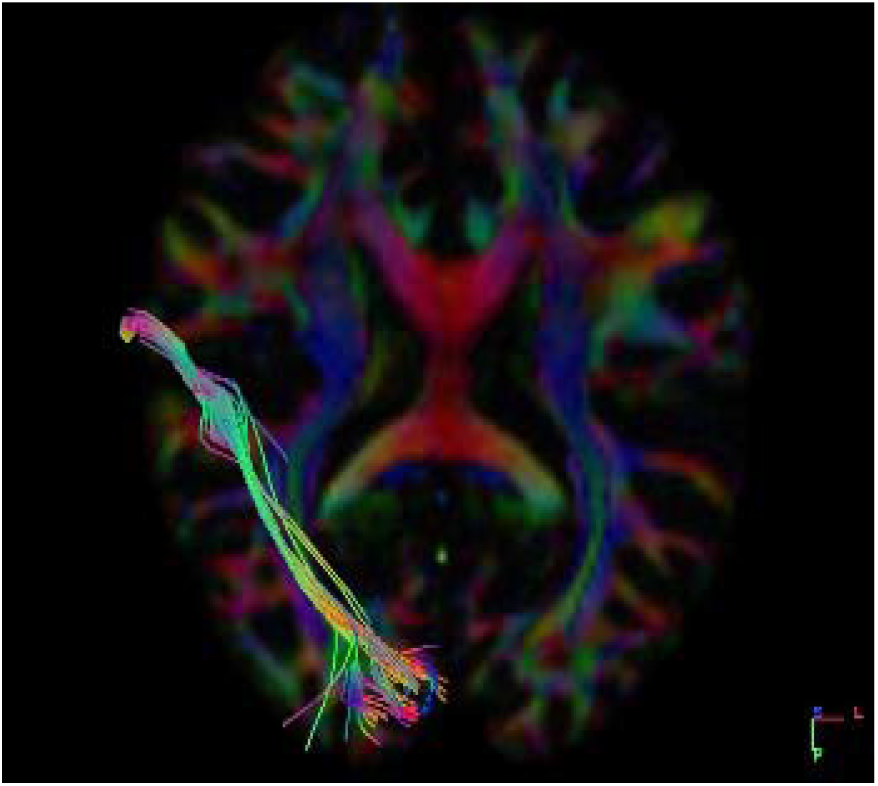
Axial Section Dataset Subject - 1024 male right side

##### Graphical Representation

**(Graph-11)** plot showing Length of tracts on x-axis and 16 subjects on y-axis to compare both left and right side of female datasets graphically. On an average, we found that the right-side has a greater Mean Length of tracts in males when compared with right side in female subjects.

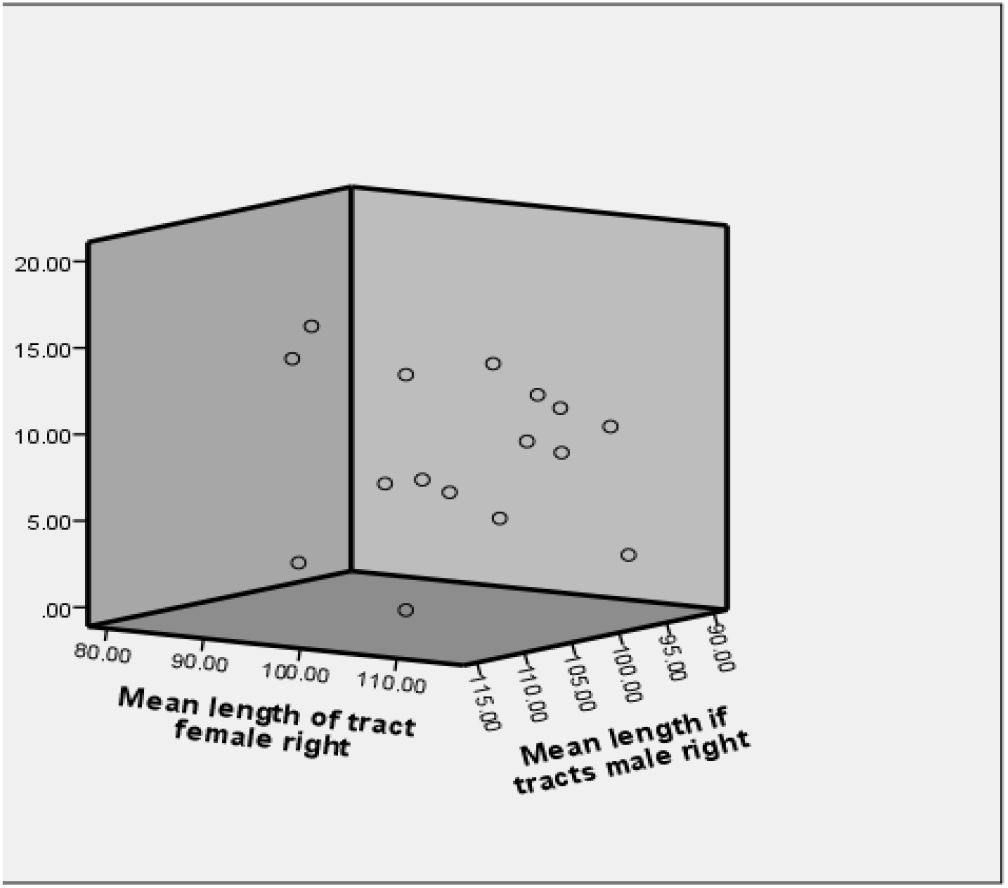

We conclude that there is no complete right-side dominance of ventral stream visual pathway
in both male and female subjects.

#### d) Left side of Male and Female subjects

The Fibres were traced and observed for neural structural connectivity on both right and left side in all sixteen healthy adult female datasets **(Table-12).**

**Table 12:** Mean length of fibres in Female and Male Left **

| S.NO | *Length of Tracts Mean(mm) Female* | *Length of Tracts Mean(mm)* |
| --- | --- | --- |
| 1 | 94.8221 | 86.2999 |
| 2 | 103.318 | 102.975 |
| 3 | 95.034 | 97.361 |
| 4 | 101.919 | 91.4074 |
| 5 | 101.059 | 108.222 |
| 6 | 94.5857 | 103.787 |
| 7 | 98.1627 | 119.844 |
| 8 | 91.3946 | 97.666 |
| 9 | 98.9259 | 88.922 |
| 10 | 91.7771 | 113.468 |
| 11 | 104.036 | 122.641 |
| 12 | 98.6967 | 119.758 |
| 13 | 100.648 | 100.107 |
| 14 | 114.255 | 109.768 |
| 15 | 104.669 | 105.942 |
| 16 | 111.022 | 105.942 |

\*\*p-value is .000031, result is significant at $p < .10$

Dataset numbers 1002, 1010, 1012, 1019, 1023, 1024, 1026, 1033, 1035 shows greater number of tracts on females. Dataset numbers 1006, 1007, 1008, 1011, 1013, 1025, 1027, 1029, 1031 shows a greater Mean Length of tracts in males. Therefore, from the table given below, 9 datasets have more fibers in male and 9 datasets have more fibers in female but the variation in the Mean Length of fibers are more seen on left side in male subject **(dataset number 1024).**

When the results were statistically compared by performing independent t–test, significant difference was seen between the Mean length of tracts in female and male subjects on left side.

**(Fig 22).**
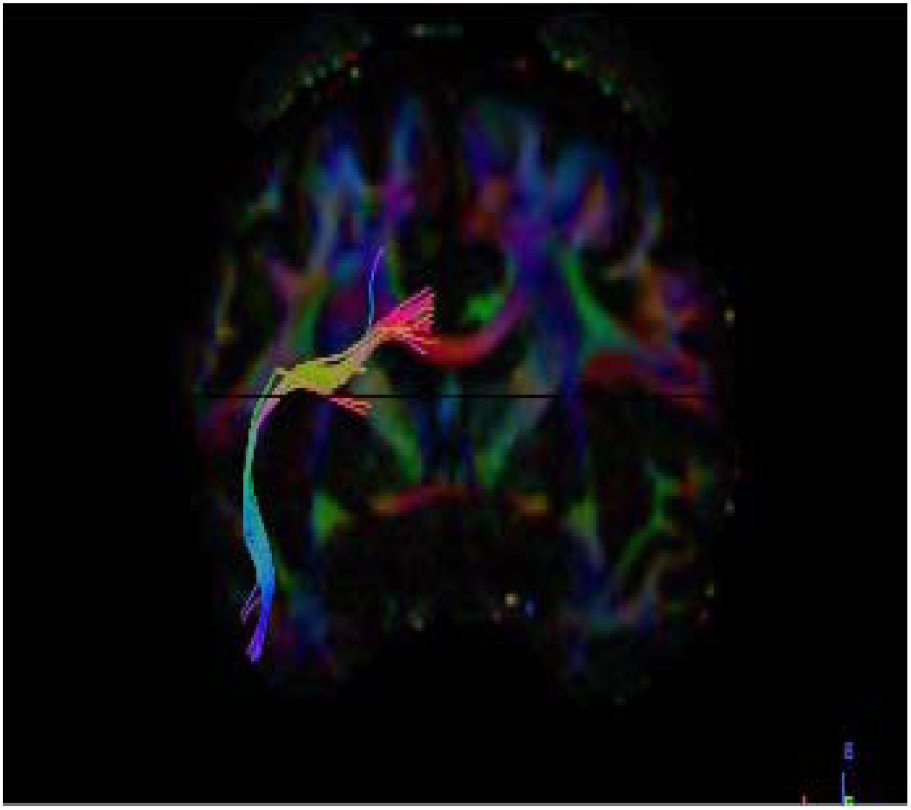
Coronal section Dataset Subject - 1024 Female left side

**(Fig 23).**
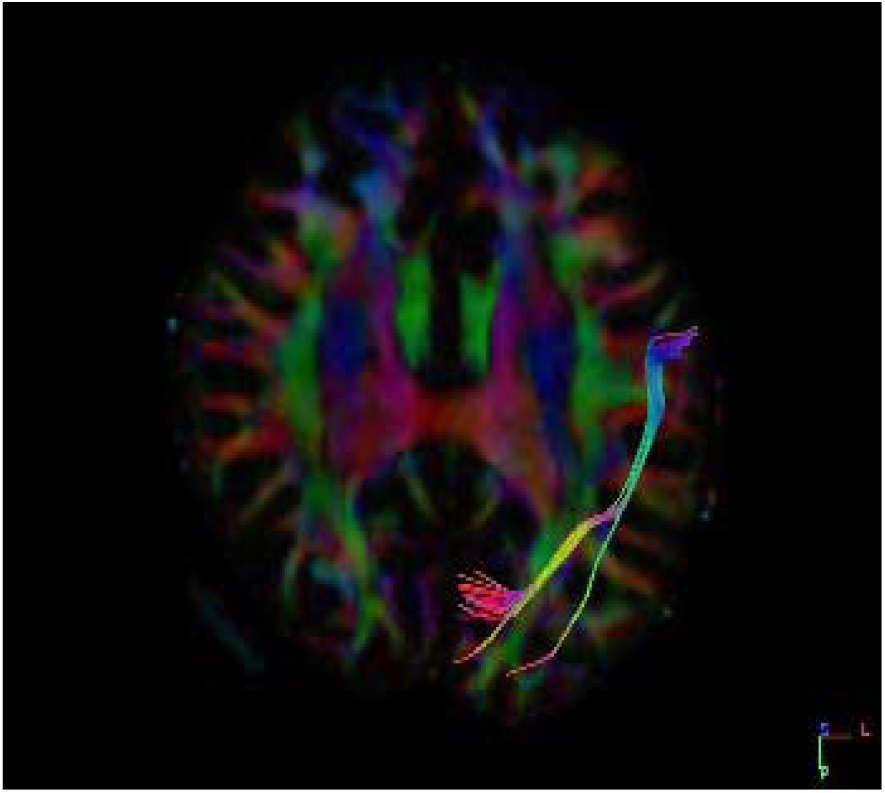
Axial Section Dataset Subject - 1024 male left side

##### Graphical Representation

**(Graph-12)** plot showing Mean Length of tracts on x-axis and **16** subjects on y-axis to compare the left of both male and female datasets graphically. On average,we found that the left-side has a greater Mean Length of tracts in males when compared with Left side in female subjects.

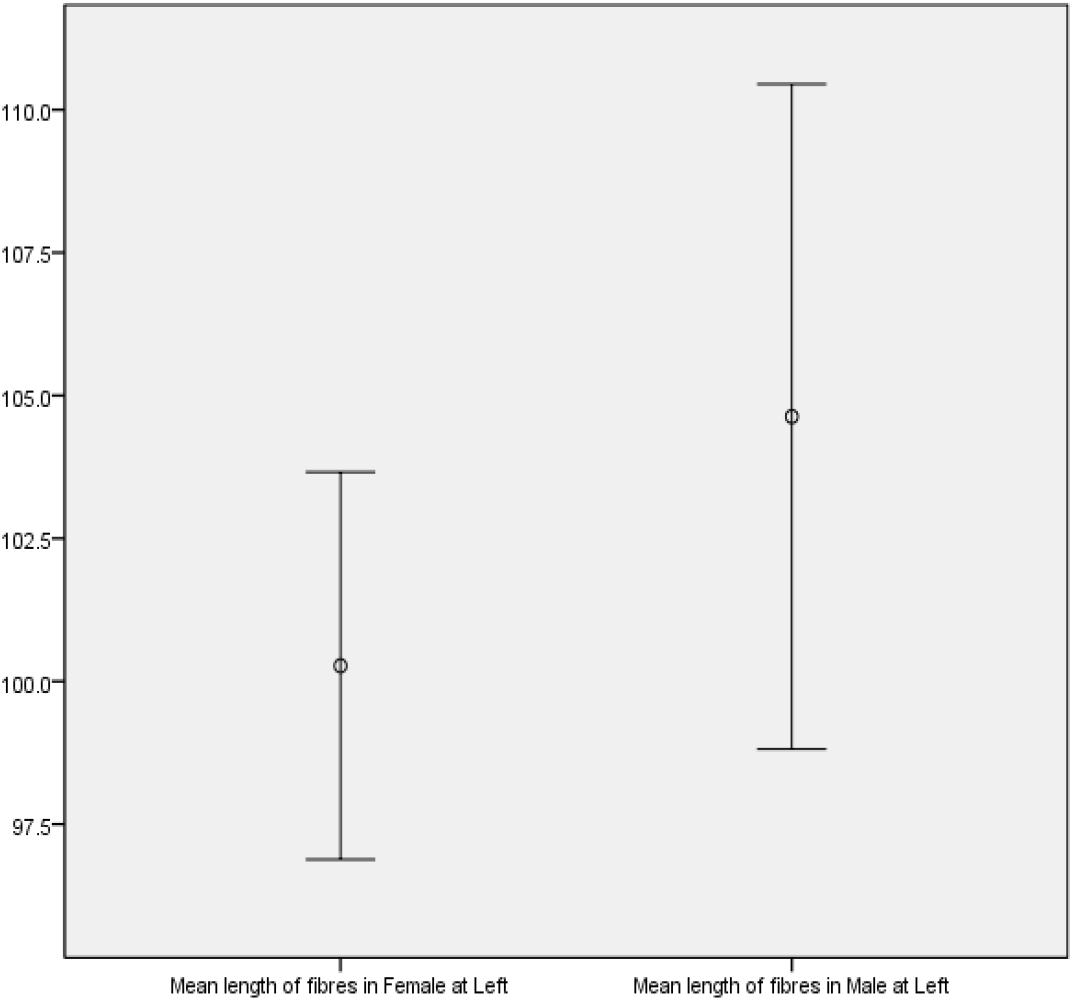

We conclude that there is complete left-side dominance of ventral stream visual pathway in both males and females. Therefore when a person has left dominance of ventral stream visual pathway, they have a better ability to interpret conceptual perception of objects.

### iv) Tract Length Standard Deviation

Tract length standard deviation is a measure that is used to quantify the amount of variation or dispersion of a set of data values. We are analysing tract mean length (mm) present in both the sexes i.e.

i)Male subjects and ii) Female subjects.

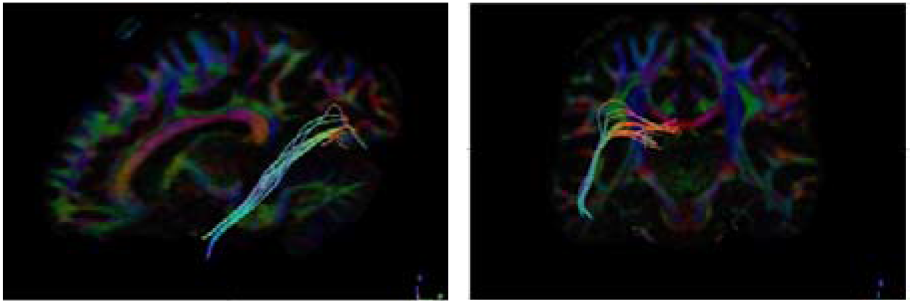

**Table 13:** TRACT LENGTH STANDARD DEVIATION IN MALE RIGHT

| **Subject** | **Gender** | **Age** | **Tracts length Mean Standard Deviation (mm<sup>3</sup>)** |
| --- | --- | --- | --- |
| 1 | Male | 40-44 | **15.9831** |
| 2 | Male | 35-39 | **5.71772** |
| 3 | Male | 25-29 | **25.004** |
| 4 | Male | 25-29 | **7.8788** |
| 5 | Male | 25-29 | **10.7874** |
| 6 | Male | 25-29 | **11.8063** |
| 7 | Male | 20-24 | **8.25238** |
| 8 | Male | 25-29 | **1.82426** |
| 9 | Male | 25-29 | **12.2629** |
| 10 | Male | 20-24 | **2.95297** |
| 11 | Male | 25-29 | **9.31404** |
| 12 | Male | 45-59 | **17.3453** |
| 13 | Male | 20-24 | **9.60575** |
| 14 | Male | 25-29 | **11819.3** |
| 15 | Male | 20-24 | **4.28517** |
| 16 | Male | 25-29 | **15.8115** |

**Table 16:**
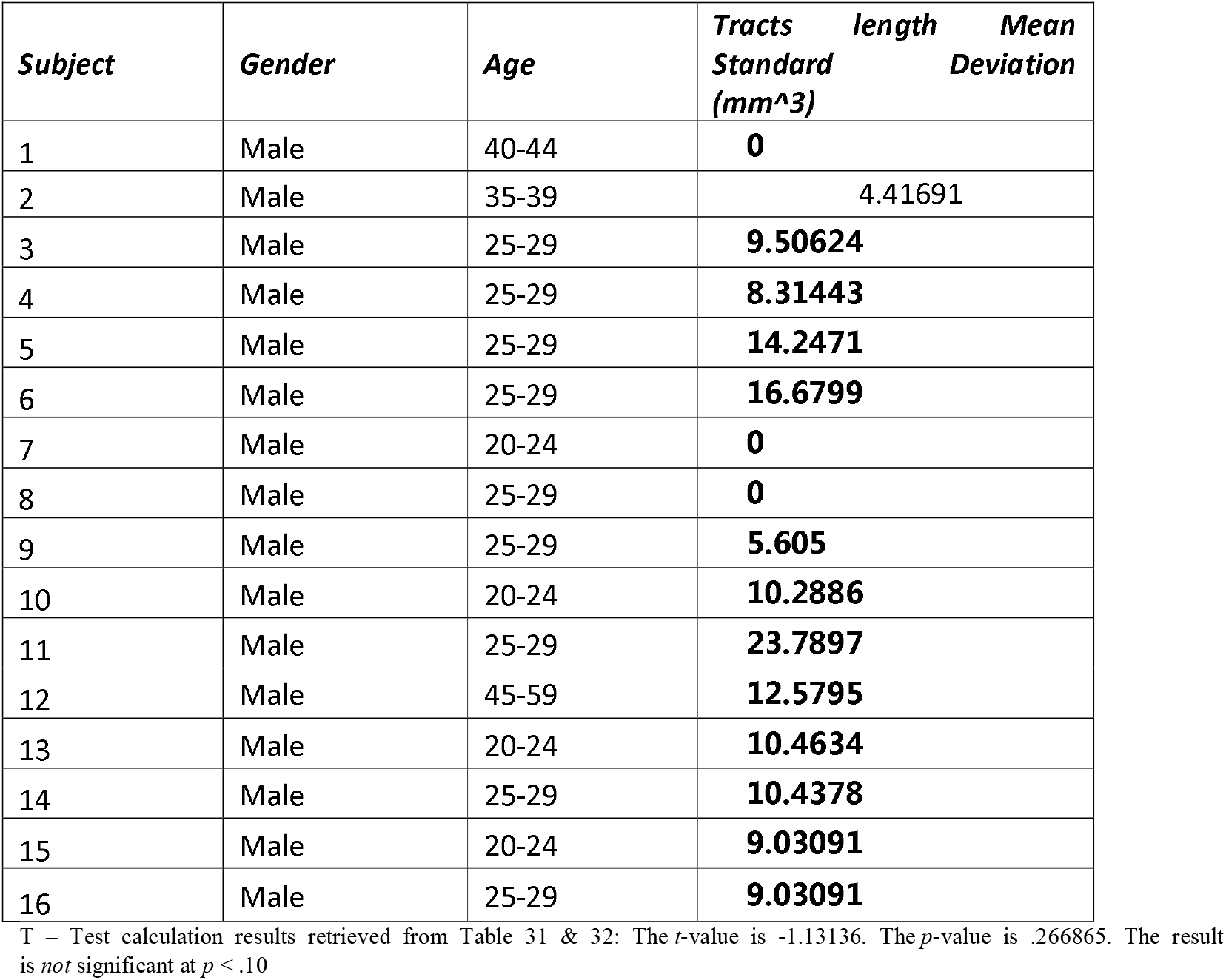
TRACT LENGTH STANDARD DEVIATION IN MALE LEFT

| **Subject** | **Gender** | **Age** | **Tracts length Mean Standard Deviation (mm<sup>3</sup>)** |
| --- | --- | --- | --- |
| 1 | Male | 40-44 | 0 |
| 2 | Male | 35-39 | 4.41691 |
| 3 | Male | 25-29 | 9.50624 |
| 4 | Male | 25-29 | 8.31443 |
| 5 | Male | 25-29 | 14.2471 |
| 6 | Male | 25-29 | 16.6799 |
| 7 | Male | 20-24 | 0 |
| 8 | Male | 25-29 | 0 |
| 9 | Male | 25-29 | 5.605 |
| 10 | Male | 20-24 | 10.2886 |
| 11 | Male | 25-29 | 23.7897 |
| 12 | Male | 45-59 | 12.5795 |
| 13 | Male | 20-24 | 10.4634 |
| 14 | Male | 25-29 | 10.4378 |
| 15 | Male | 20-24 | 9.03091 |
| 16 | Male | 25-29 | 9.03091 |

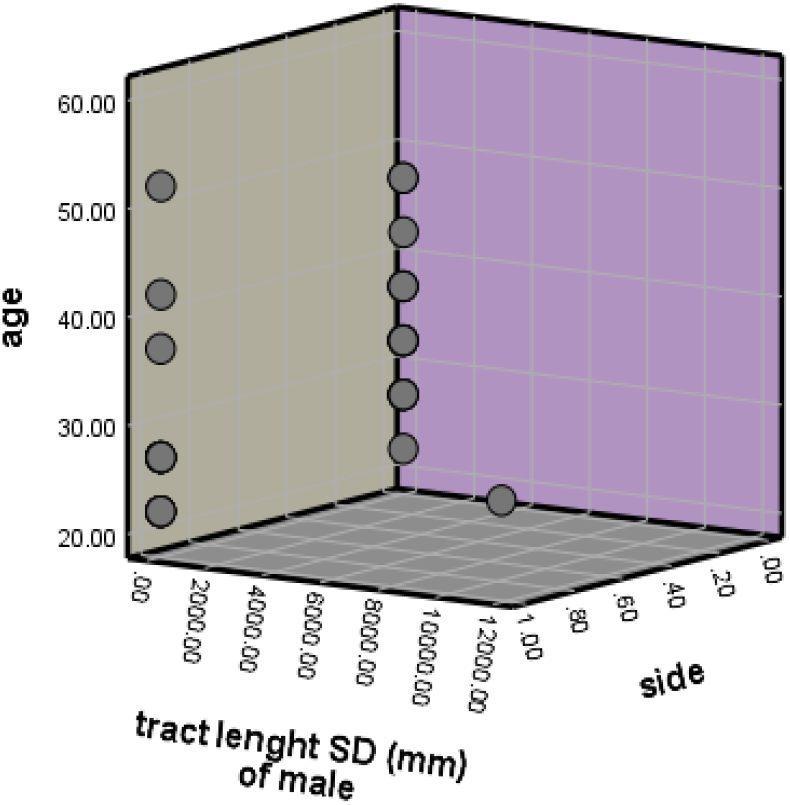

**Table 17:**
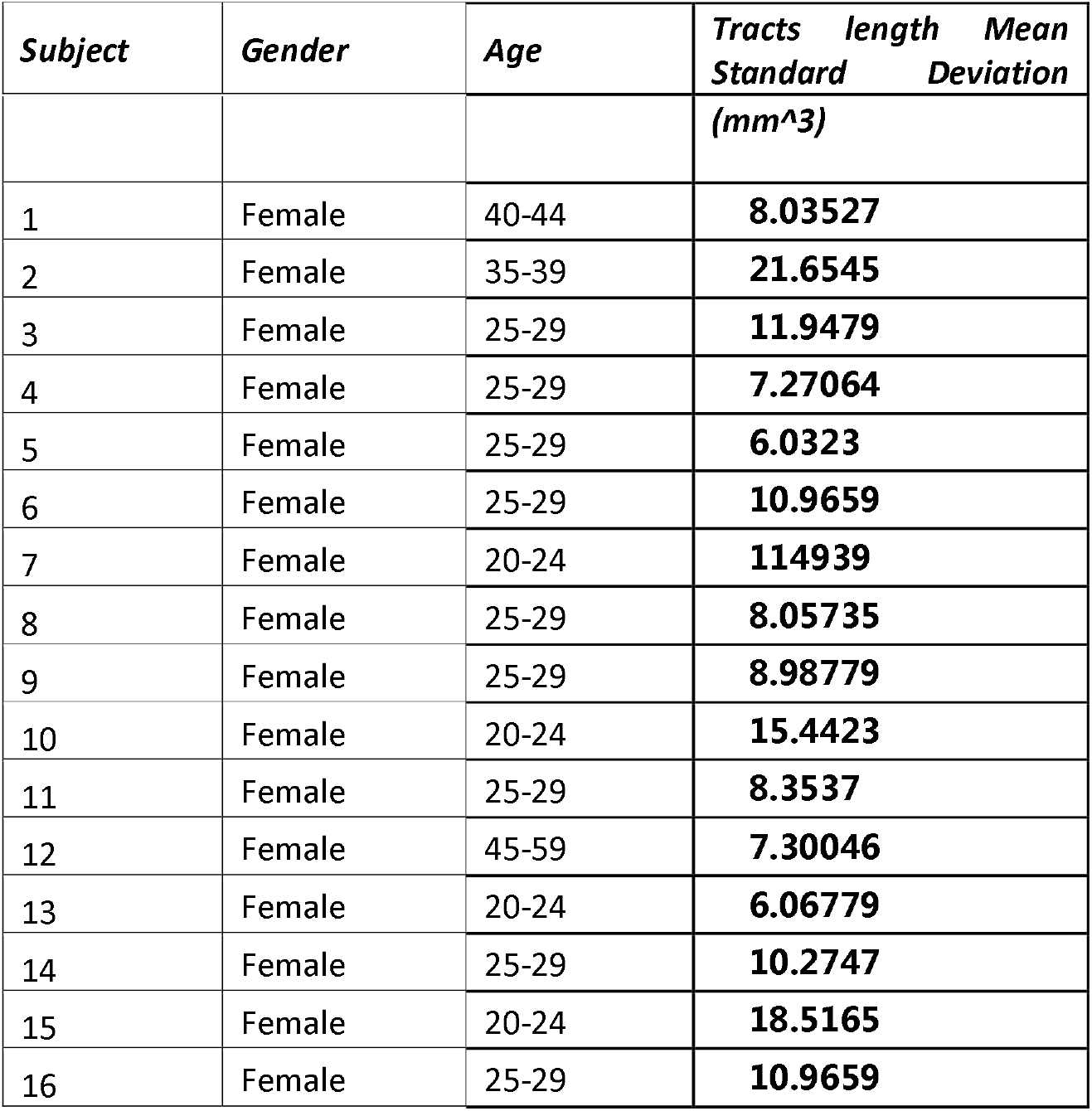
TRACT LENGTH STANDARD DEVIATION IN FEMALE RIGHT

**Table 18:** TRACT LENGTH STANDARD DEVIATION IN FEMALE LEFT

| **Subject** | **Gender** | **Age** | **Tracts length Mean**<br>**Standard Deviation**<br>**(mm<sup>3</sup>)** |
| --- | --- | --- | --- |
| 1 | Female | 40-44 | **7.03841** |
| 2 | Female | 35-39 | **2.65183** |
| 3 | Female | 25-29 | **11.7095** |
| 4 | Female | 25-29 | **4.0939** |
| 5 | Female | 25-29 | **16.7269** |
| 6 | Female | 25-29 | **7.97714** |
| 7 | Female | 20-24 | **8.31999** |
| 8 | Female | 25-29 | **5.75632** |
| 9 | Female | 25-29 | **8.86888** |
| 10 | Female | 20-24 | **6.33233** |
| 11 | Female | 25-29 | **12.3925** |
| 12 | Female | 45-59 | **12.2787** |
| 13 | Female | 20-24 | **12.3416** |
| 14 | Female | 25-29 | **8.73704** |
| 15 | Female | 20-24 | **8.91893** |
| 16 | Female | 25-29 | **13.1323** |

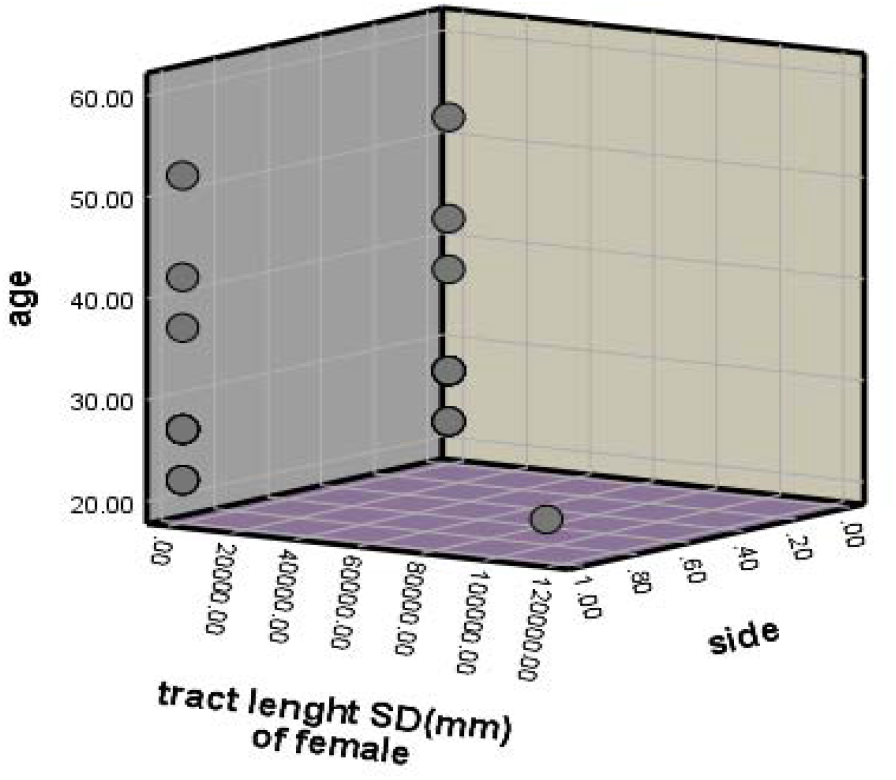

## 4. Discussion and Conclusion

According to previous records and publications, there are many well-known scientists who have spent their lives to study the mechanism of dorsal and ventral stream of the vision. The great finding of two separate mechanisms of visual systems in monkey was stated by Treva in 1968 (1). This formed a path for the establishment and identification of dorsal and ventral stream. In 1969, Schneider stated the two visual pathways for locating and identifying objects (2). Later, Ingle in 1973 visualized the two systems in frogs (3). After 9 years, i.e. in 1992, Ungerleider and Mishkin were able to differentiate the dorsal and ventral streams with respect to processing spatial and visual features, based on studies and research done on monkeys (4). But it wasn’t a long success as it was superseded by Milner & Goodale in later years. The existence of functional differences in the dorsal and ventral streams were stated by Huxby in 1994 by proposing the “What” and “Where” as two different pathways through positron emission tomography (5). In 2002, Norman stated the two theories of perception, where the ventral streams’ central function is identification of objects (6). Polanen and Davere in 2015 stated that there is an interaction between the dorsal and ventral streams where the dorsal is fed by the ventral. They also proposed that there is a direct white matter connection between dorsal and ventral which helps them to interact each other bidirectionally. Hence, scientists previously haven’t given any proof of structural connection, however, we are postulating a new concept of structural connection where the connectivity of the ventral stream has been stated. In this article, we have proposed that the ventral stream visual pathway has a neuronal structural connectivity where a group of fibres in the central nervous system forms tracts that pass from the primary visual cortex to the inferior temporal lobe and enables individuals to perceive objects. Additionally, the fibres that interconnects Brodmann area 17,18,19 to Brodmann area 20 allows for flow the visual information for the perception of an object thus, forming an occipitotemporal network. This interesting connection forming a true occipitotemporal network is what helps a person perceive an object, its shape, colour and also the different characteristics of the object.

Hence, during our study, we analyzed a list of parameters inclusive of number of tracts, tract length, its volume and standard deviation of the tract length which could be useful in recognizing the cause for different neurodegenerative disorders. Typical long-term memory and the ability to organize an object spatially are the most important function of ventral stream visual pathway. While the dorsal helps the ventral stream to interpret the object by sending the spatial information of dorsal to the ventral stream. But the major question is how both the dorsal and ventral streams coordinate structurally with each other to provide a pure output of information about the object location. Analysis from our study shows that females and males predominately possess similar number of fibres passing from the primary visual cortex to the inferior temporal lobe. When it comes to the tract volume and tract mean standard deviation (SD) of the fibres, there was a vast difference seen with females having more volume and SD than the males. The tract length of the fibres in both male and female showed similar result.. Hence, we conclude by stating that the there is an existence of a structural connection enabling the transfer of information. Polanen mentioned that bilateral damage to the inferior temporal lobe leads to damage to the ventral stream visual pathway and and this deprived some patients of the ability of object perception, hence, any damage to the ventral stream may lead the patients developing apperceptive visual agnosia. Apperceptive form of visual agnosia is where patients suffer lack of coordination and they do not usually recognize objects and hence cannot grasp objects thus, this shows there is a lack of interaction of the ventral stream with the dorsal stream.

